# Structural covariance of brain region volumes is associated with both structural connectivity and transcriptomic similarity

**DOI:** 10.1101/183004

**Authors:** Yohan Yee, Darren J. Fernandes, Leon French, Jacob Ellegood, Lindsay S. Cahill, Dulcie A. Vousden, Leigh Spencer Noakes, Jan Scholz, Matthijs C. van Eede, Brian J. Nieman, John G. Sled, Jason P. Lerch

## Abstract

An organizational pattern seen in the brain, termed *structural covariance*, is the statistical association of pairs of brain regions in their anatomical properties. These associations, measured across a population as covariances or correlations usually in cortical thickness or volume, are thought to reflect genetic and environmental underpinnings.

Here, we examine the biological basis of structural volume covariance in the mouse brain. We first examined large scale associations between brain region volumes using an atlas-based approach that parcellated the entire mouse brain into 318 regions over which correlations in volume were assessed, for volumes obtained from 153 mouse brain images via high-resolution MRI. We then used a seed-based approach and determined, for 108 different seed regions across the brain and using mouse gene expression and connectivity data from the Allen Institute for Brain Science, the variation in structural covariance data that could be explained by distance to seed, transcriptomic similarity to seed, and connectivity to seed.

We found that overall, correlations in structure volumes hierarchically clustered into distinct anatomical systems, similar to findings from other studies and similar to other types of networks in the brain, including structural connectivity and transcriptomic similarity networks. Across seeds, this structural covariance was significantly explained by distance (17% of the variation, up to a maximum of 49% for structural covariance to the visceral area of the cortex), transcriptomic similarity (13% of the variation, up to maximum of 28% for structural covariance to the primary visual area) and connectivity (15% of the variation, up to a maximum of 36% for structural covariance to the intermediate reticular nucleus in the medulla) of covarying structures. Together, distance, connectivity, and transcriptomic similarity explained 37% of structural covariance, up to a maximum of 63% for structural covariance to the visceral area. Additionally, this pattern of explained variation differed spatially across the brain, with transcriptomic similarity playing a larger role in the cortex than subcortex, while connectivity explains structural covariance best in parts of the cortex, midbrain, and hindbrain. These results suggest that both gene expression and connectivity underlie structural volume covariance, albeit to different extents depending on brain region, and this relationship is modulated by distance.

## 1. Introduction

Patterns of covariation in the thickness or volume of brain regions (“structural co-variance”), measured across a population, have been linked to both structural and functional networks of the brain. Previously, Gong et al. (2012) showed that approximately 35-40% of cortical regions that positively correlated in thickness were also connected by fibre tracts estimated from probabilistic tractography on diffusion MRI data. The spatially widely-distributed nature of structural covariance networks suggest that they might arise from functional connectivity along with specific fibre connections; Lerch et al. (2006) demonstrate that cortical thickness covariance arises between structurally and functionally connected regions, and Segall et al. (2012) provide evidence that functional connectivity might also explain structural covariance (of gray matter density) by showing prominent correlations between many independent component pairs of structural covariance and resting state networks. More recently, Reid et al. (2016) use cross-species data to show a correspondence between cortical thickness networks, tractographic networks obtained from diffusion-weighted MRI (DWI), and resting-state fMRI; here, approximately 15% of cortical thickness covariance was predicted by DWI and fMRI in humans, and 25% in macaques. Together, these studies point to a link between connectivity and structural association of brain regions. Indeed, given this link to connectivity, structural covariance networks are particularly appealing to examine neuropsychiatric disorders in which aberrations in structural and functional networks have been implicated. Alterations in networks of structural covariance have been demonstrated in autism (Zielinski et al., 2012; Bernhardt et al., 2014; Valk et al., 2015; Bethlehem et al., 2017), schizophrenia (Shi et al., 2012; Wheeler et al., 2014; Alexander-Bloch et al., 2014), epilepsy (Bernhardt et al., 2011; Yasuda et al., 2015; Bernhardt et al., 2016), and grapheme-color synesthesia (Hänggi et al., 2011), to name a few such disorders.

The mechanisms that underlie structural covariance have yet to be well characterized. Correlations with structural and functional networks suggest that structural covariance might arise due to network mediated plasticity—regions that fire together and wire together might also couple in volumes together due to mutually trophic, plasticity-related changes at the synaptic and cellular levels (Evans, 2013). The previous studies mentioned suggest that this plasticity might only partially account for structural covariance. While it is likely that this might be due to methodological constraints (for example, estimates of the proportion of white matter voxels which contain crossing fibres range from a third (Behrens et al., 2007) to 90% (Jeurissen et al., 2013), making comparisons to tractography-estimated structural connectivity challenging), other biological factors might also explain covariation patterns. Another such (not necessarily mutally exclusive) mechanism is coordinated neurodevelopment (Alexander-Bloch et al., 2013a; Evans, 2013). Alexander-Bloch et al. (2013b) showed that networks of cortical thickness covariance agree strongly with networks of cortical thickness *change*, a measure of this synchronized neurodevelopment. Such networks of anatomical change (“maturational coupling”) are conjectured to arise from the expression of common genetic cues during early development of the cortex (Raznahan et al., 2011). Supporting this are twin studies implicating genetics and structure (Schmitt et al., 2008; Rimol et al., 2010; Docherty et al., 2015), with one by Schmitt et al. (2008) suggesting that the small-world network organization of structural covariance (He et al., 2007) might be explained by genetic correlations that display a similar pattern. The extent that transcriptomic similarity mediates covariance, particularly in relation to connectivity, remains to be seen, however. Nevertheless, given this link between neurodevelopment, genetics, and structural covariance, it is not surprising that alterations in structural covariance arise in relation with aberrant gene expression (Pezawas et al., 2008; Schmitt et al., 2016; Bruno et al., 2016) or early sensory deprivation (Voss & Zatorre, 2015).

To probe the mechanisms that underlie structural covariance and examine the role of genetics and connectivity in particular, we asked the question, *to what extent do transcriptomic similarity and structural connectivity underlie structural volume covariance?* Here, we leveraged connectivity and gene expression data from the Allen Institute for Brain Science in order to address this question in the mouse brain. Genetic and environmental control of mice allow for the comparison of structural covariance to connectivity and expression similarity in highly similar populations. Pagani et al. (2016) have shown that networks of structures that covary together in volume, consistent with neuroanatomical systems, emerge in an analysis of structural covariance in the mouse brain. A seed-based approach further shows the presence of bilateral and neuroanatomically specific networks of covariance (Pagani et al., 2016). In this study, we first analyze parcellation-derived networks constructed from MR images of mouse brains in relation to connectivity, transcriptomic similarity networks, and distance between structures. Then, using a seed-based approach with 108 injection sites from the Allen Institute’s mouse connectivity experiments as seeds, we examine the variation in structural covariance that can be explained by transcriptomic similarity, structural connectivity, and physical distance to seed, and explore the spatial pattern of this explained variation.

## 2. Methods

### 2.1. Outline and definitions

In this study, we use the term *structural covariance* to describe correlations in volumes between pairs of regions. We examine the biological basis of structural covariance in two separate ways: 1) using a parcellation-based approach in which structural covariance is computed between the volumes of regions that are defined by a 318 structure neuroanatomical atlas, and 2) a seed-based in which structural covariance is computed for the whole brain in a voxelwise manner to each seed, for a set of 108 seed regions. In both cases, we used the Pearson correlation coefficient as a measure of structural covariance because, unlike the unscaled covariance, it does not span many orders of magnitude.

In the parcellation-based approach, we examine the spatial structure of the structural covariance (correlation) matrix, and compare this to similarly constructed matrices for transcriptomic similarity, structural connectivity, and Euclidean distance.

In the seed-based approach, we examine the variation in structural covariance values at each voxel (i.e. correlation coefficients) that can be explained by transcriptomic similarity, structural connectivity, and Euclidean distance. To do so, we construct a structural covariance map (i.e. a 3D dataset) for each of the 108 seed regions and fit linear models with structural connectivity, transcriptomic similarity, and distance to seed as predictors for structural covariance values. For a given structural covariance map and a model, we used the *R*^2^ value (adjusted for multiple predictors where applicable) of the linear model to quantify the extent that a structural covariance map is associated with the model’s predictors; this is the variation in the structural covariance data that can be explained by the model (*variation explained* for short).

### 2.2. External data sources

For this study, we used mouse connectivity (Oh et al., 2014) and gene expression (Lein et al., 2007) data from the Allen Institute for Brain Science. The mouse connectivity data consists of neuronal tracers injected into a variety of regions in the mouse brain that show projections that emanate from the injection sites. Neuronal tracers avoid tractography-related issues that arise when inferring connectivity from diffusion MRI data, and allow for the visualization of fine tracts that might not be detected through MRI. The mouse gene expression data consists of 3D images for a set of genes that show the spatial expression pattern of each gene, and is the most comprehensive high-resolution dataset to date.

#### Mouse connectivity

Data from the Allen Institute’s mouse connectivity experiments (Oh et al., 2014) were used to assess structural connectivity between structures. In a series of tracer injection experiments, the Allen Institute injected a recombinant adeno-associated viral (rAAV) tracer that expresses enhanced green fluorescent protein (EGFP) under control of a human synapsin I promoter and thereby labels neurons. Injections (for the data used in this study) were in adult (age: postnatal days P56 ± 2) male wildtype C57Bl6/J mice. The tracer used does not cross synapses to label further downstream axons, and thus describes directed, monosynaptic connectivity. The injected brains were imaged by the Allen Institute for Brain Science using serial two-photon microscopy at an in-slice resolution of 0.35*μ*m and coronal slice interval of 100*μ*m, and further processed. Processing steps included intensity correction and stitching of images, followed by a nonlinear alignment to a 3D reference model that forms the basis of the Allen Institute defined Common Coordinate Framework (Version 3, “CCFv3”). Further processing to detect EGFP expression includes intensity rescaling, noise removal, tissue segmentation, and projection signal segmentation. As a summary of the high-resolution projection data, the Allen Institute made available the *projection density*, a 3D image that grids the post-processed fluorescence data at 50 *μ*m and expresses the proportion of voxels at original resolution which show a tracer signal. These projection density (50 *μ*m grid) data, which consists of a 3D image ranging in values between 0 and 1 (inclusive) per injection site, describes anterograde connectivity from the injection site. All projection density data (50 *μ*m grid aligned in CCFv3 space) for injections in wildtype C57Bl/6J mice consisting of a total of 488 injection experiments^1^ were downloaded.

#### Mouse gene expression

To assess transcriptomic similarity, 4345 3D gene expression images for 4082 unique genes were downloaded from the Allen Institute’s coronal expression dataset (Lein et al., 2007) were used. Gene expression data were obtained by the Allen Institute following a pipeline that involved semi-automated riboprobe generation, tissue preparation and sectioning, *in-situ* hybridization (ISH), imaging, and data post-processing. Mice used for these gene expression studies were similar in age (postnatal day P56), sex (male), and strain (C57Bl/6J) to those used in the connectivity studies. Briefly, for a given expression image, a mouse brain was sectioned into 8 series of slices, 6 of which were hybridized to the given gene and 2 of which were Nissl stained for anatomical reference. The Nissl and expression images obtained from the ISH experiments were processed by the Allen Institute in steps that included intensity and white balance normalization, separation of foreground from background, removal of noise, connected component analysis, alignment to a 3D reference model, tissue segmentation, and expression detection. The Allen Institute provided summaries of the spatial expression data at a 200 *μ*m resolution, termed the *gene expression energy*. This gene expression energy, defined as the sum of expressing pixel intensity divided by the sum of all pixels in a division, increases in regions of high expression, and is bounded by zero in regions of no expression. Note that the Allen Institute also provides a sagittal expression dataset comprising of images for ~20000 genes. We chose to use the coronal dataset because of its whole-brain coverage and quality.

### 2.3. Animals and imaging

Structural covariance is a property of a population and is therefore measured over a group of individuals. Here, we constructed structural covariance networks in a group of 153 mice imaged via MRI. Ex-vivo images were high resolution and covered the whole brain.

MR images were obtained in-house at the Mouse Imaging Centre in a multiple mouse imaging setup (Lerch et al., 2011a) and as part of other studies’ wildtype groups (Ellegood et al., 2015; Cahill et al., 2015). All 153 images were T2-weighted and obtained ex-vivo on a 7T Varian MR scanner, with brains perfused with a gadolinium-based contrast agent before imaging (de Guzman et al., 2016). Since the images were collected over a period of several years, a variety of MR scan parameters were used to obtain the images, which ranged in resolution from 32-56*μ*m (isotropic). Mice were selected to match those used in the Allen Institute for Brain Science’s connectivity and gene expression experiments in terms of strain, sex, and age (adulthood). As such, mice were male C57Bl/6 and adults (ranging in age from postnatal days P60-112). Some mice underwent interventions (exercise wheel in cage, saline injection). Table 1 describes all mice used.

**Table 1:** Mouse data and imaging parameters. All ages refer to the age in postnatal days at which the mice were perfused. Source acronyms refer to author initials. T2w FSE stands for T2-weighted fast spin echo. All resolutions reported are isotropic.

| Study | Mouse data |  |  |  | Imaging parameters |  |  |  |  |
| --- | --- | --- | --- | --- | --- | --- | --- | --- | --- |
| Sample size | Strain | Sex | Age (days) | Intervention | Source | Scanner | Sequence | Resolution | Data collected |
| 8 | C57Bl/6 | M | 64 | Saline injection | LSN | Varian 7T | T2w FSE | 40μm | 2015 |
| 10 | C57Bl/6 | M | 60 | Saline injection | JE | Varian 7T | T2w FSE | 56μm | 2012-2015 |
| 10 | C57Bl/6 | M | 60 | Saline injection + maternal separation | JE | Varian 7T | T2w FSE | 56μm | 2012-2015 |
| 22 | C57Bl/6 | M | 60 | None | JE | Varian 7T | T2w FSE | 56μm | 2012-2015 |
| 11 | C57Bl/6 | M | 60 | None | JE | Varian 7T | T2w FSE | 56μm | 2012 |
| 9 | C57Bl/6 | M | 60 | None | JE | Varian 7T | T2w FSE | 32μm | 2010 |
| 10 | C57Bl/6 | M | 60 | None | JE | Varian 7T | T2w FSE | 40μm | 2013 |
| 13 | C57Bl/6 | M | 60 | None | JE | Varian 7T | T2w FSE | 56μm | 2011-2013 |
| 30 | C57Bl/6 | M | 112 | None | LSC | Varian 7T | T2w FSE | 56μm | 2012 |
| 30 | C57Bl/6 | M | 112 | Exercise wheel | LSC | Varian 7T | T2w FSE | 56μm | 2012 |

### 2.4. Registration and volumes

Deformation-based morphometry was used to register the mouse brains images (after correcting for geometric distortions) to a common non-linear average brain (Lerch et al., 2011a). The purpose of registration was to determine volumes of neuroanatomical regions required to compute the structural covariance networks. The 153 images were registered in four separate groups based on the images’ experimental source and environmental interactions; images were registered to group consensus averages in an iterative pipeline (Lerch et al., 2011a). Registering to separate group averages is analogous to regressing out volume differences resulting from different exposures. Nonlinear registration was achieved using ANTS (Avants et al., 2009). The PydPiper framework (Friedel et al., 2014) was used for image registration; registration was carried out on the General Purpose Cluster at the SciNet HPC Consortium (Loken et al., 2010). The registration procedure outputs a series of spatial transformations that map the non-linear average of all images to each input image, along with a corresponding set of Jacobian determinant images that are measures of local volume deviations of each mouse from the average image. The Jacobian determinant images were further log-transformed to reduced skewness (Leow et al., 2007). Structure volumes within each mouse image were computed by summing over Jacobian determinants at each voxel within the structure as defined by the atlas after mapping onto the average image (Lerch et al., 2011a).

All analyses were carried out in CCFv3 space. Nonlinear average images of each group were registered individually to the two-photon microscopy CCFv3 (50 *μ*m in-slice resolution) reference average from the Allen Institute using ANTS. Invidiual images (including the Jacobian determinant images) were further transformed to CCFv3 space on the basis of the transformation defined between the average and CCFv3 space, and then resampled at 50 *μ*m isotropic resolution, thereby allowing direct voxelwise comparisons across all images.

Using an atlas which defines 318 structures (see Section 2.5) that cover the whole brain, structure volumes were computed for each mouse allowing for an atlas-based exploration of structural covariance. The seed-based analyses was carried out voxelwise, using log-transformed Jacobian determinants as a measure of local volumes.

### 2.5. Parcellation-based exploration

To explore large-scale patterns of structural covariance in the mouse brain, an atlas-based approach was used in which the correlations between the volumes of predefined brain structures were computed.

We used an atlas that defines 318 structures in total when considering bilateral structures separately; this atlas combined a high-resolution three-dimensional brain atlas of C57Bl/6J mice by Dorr et al. (2008) with a segmentation of cerebellar structures by Steadman et al. (2014) and a segmentation of the neocortex by Ullmann et al. (2013) (the “Dorr-Steadman-Ullmann” or DSU-atlas). To avoid spurious correlations driven by whole-brain volume, we considered for each mouse the normalized volumes of these structures, relative to the whole-brain volume (i.e. percent volume).

For each pair of the 318 DSU-atlas regions, the Pearson correlation coefficient was computed between relative structure volumes and over all individual mouse brain images, resulting in a 318x318 matrix of correlations representing the group-wise structural volume covariance network. Given that the Allen Institute’s mouse connectivity experiments consisted of injections only in the right hemisphere, the structural covariance matrix was subsetted to include only source structures in the right hemisphere (target structures in both the right/ipsi-and left/contralateral hemispheres were kept).

#### Structural connectivity matrix

We used correlations in tracer fluorescence as a measure of connectivity between structures in the parcellation-based analysis. For each projection density image, tracer projection density values from the Allen Institute were averaged over voxels in each of the 318 structures. Correlations in average tracer projection density values were computed between every pair of regions, and over a set of tracer projection density images. The set of tracer images used included all projection density images from the 488 injection experiments, along with the same 488 images flipped across the mid-sagittal plane to account for contralateral afferent connectivity. For the parcellation-based analysis, we used correlations over tracer images as a metric of connectivity rather than the raw projection density values since this describes bidirectional connectivity (efferent and afferent) via a symmetric matrix, is the same measure of association as volume correlations and transcriptomic similarity (and does not scale across multiple orders of magnitude), and allows for visually clear comparison of clusters. Directional information is maintained in the seed-based analysis below (Section 2.6).

#### Transcriptomic similarity matrix

Mean gene expression energies were computed within each of the 318 DSU-atlas defined regions for each gene. This was done by downsampling the DSU-atlas labels at the 200 *μ*m resolution of the expression images, and then by averaging each gene’s expression energy values within each region (thus providing a 4345 х 318 table). For a given gene (row), values were further normalized by dividing each element by the total mean expression (i.e., row sum). A correlation matrix representing transcriptomic similarity was computed by correlating pairwise these normalized mean expression of 4345 genes under each pair of structure labels. Expression images were not processed any further to remove any noise or missing data artefacts since these were rare within any given structure, and this noise was expected to be overcome by the strong correlation signal driven by large sample size.

#### Distance matrix

Pairwise distances were computed between all pairs of 318 structures as the Euclidean distance between structure centroids.

#### Matrix comparisons and statistical methods

The structural covariance matrix data were clustered to determine which regions form communities of similar interregional correlations. Specifically, correlations in volume between each source structure and all target structures were represented as a vector. These vectors were hierarchically clustered (using average linkage) to determine structures that tend to associate together in structural covariance patterns. The optimal number of clusters was determined by examining using a *scree* plot in which the within-sum-of-squares (WSS) cluster distance is plotted for different cluster numbers; the optimal cluster number is taken to be the cluster number above which an increase in the number of clusters results in little change in the WSS.

Apart from visual comparisons, a partial least-squares (PLS) analysis was used to quantify the correspondence between the visually similar structural covariance and transcriptomic similarity matrices. In this analysis, structural covariance and transcriptomic similarity matrices, subsetted to include regions in the right hemisphere, were decomposed to maximize the covariance between component matrices.

### 2.6. Seed-based voxelwise analysis

In addition to the parcellation-based analysis described above, we used a seed-based approach to examine the relationship between structural covariance and physical distance, transcriptomic similarity, and structural connectivity. In this approach, we constructed structural covariance maps voxelwise to predefined seed regions of interest. Our approach was to examine the variation in these structural covariance data that could be attributed to a) neuronal tracer data from the Allen Institute, b) transcriptomic similarity images constructed from Allen Institute expression data, and c) physical distance to the seed. As described below, seed regions were selected from the Allen Mouse Brain Connectivity Atlas injection sites.

#### Seed selection criteria

The 488 injection experiments (in wildtype C57Bl/6J mice) from the Allen Institute’s mouse connectivity dataset (Oh et al., 2014) provided a corresponding set of injections sites, which we considered as the seed regions of interest. We found that tracer tract volume (i.e. volume of voxels outlined by tracer) and projection length depended on the volume of the injected tracer when the amount of tracer injected was small, suggesting that in this volume regime, projection tracts might be missed out when not enough tracer was injected. We thus selected only connectivity experiments in which the injection volume was >0.4 mm^3^ as reported by the Allen Institute; above this threshold the dependence of tract volume and length on injection volume was not apparent (see Figure 1a,b). 108 injection sites (51 in the cortex [Allen Institute classification: “cerebrum”], 57 in the subcortex [Allen Institute classification: “brain stem”]) matched this criterion and were considered as seed regions for this study. No cerebellar or olfactory bulb seed regions matched this criterion. Figure 1c shows the spatial distribution of these 108 seed regions, which cover approximately 18% of grey matter in the right hemisphere.

**Figure 1:**
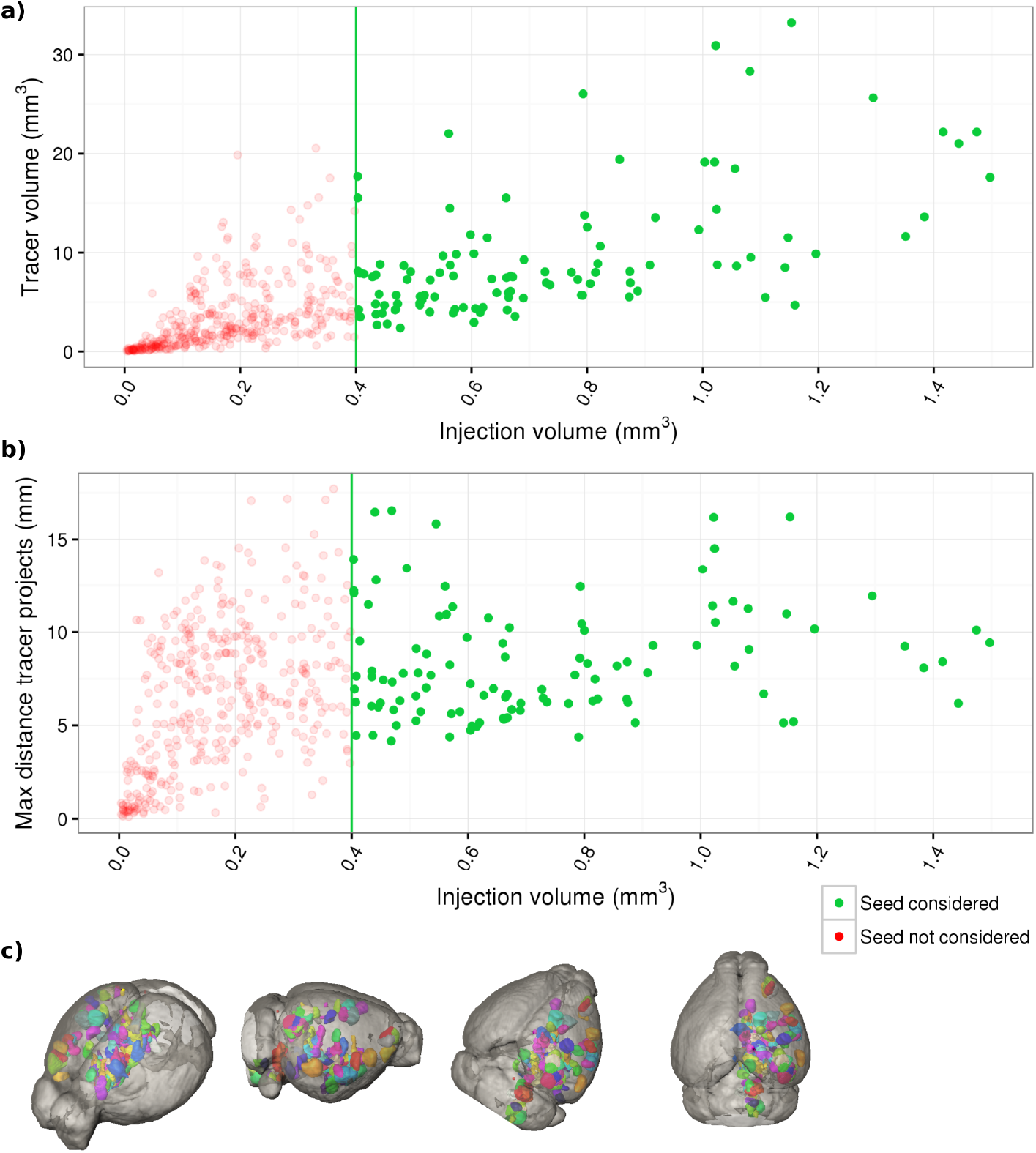
Injection experiments in which the volume of the injection was greater than 0.4 mm^3^ were considered in order to avoid experiments where projection tracts are missed due to not enough tracer uptake. Above this threshold, **(a)** the volume of voxels which show a tracer signal and **(b)** the maximum distance the tracer projects both do not depend on the injection volume (compared to the strong relationship below the threshold). A total of 108 seed regions fit this constraint; renderings of these seed (injected) regions in the mouse brain were considered for this study are shown in **(c)**. Coverage of injection regions is hemisphere-wide, with the notable exceptions of the cerebellum and olfactory bulbs.

#### Connectivity 3D datasets

We used the projection density values associated with each seed for the voxelwise analysis. These projection density data, aligned to the structural covariance data, allows for direct voxelwise comparisons between the two datasets.

#### Estimated polysynaptic connectivity 3D datasets

The rAAV tracer used in generating the connectivity datasets does not cross the synapse. We generated a prediction of what the tracer image would look like if the tracer could “hop” across synapses by combining overlapping tracer images; Figure S2 is an illustrative example of this procedure.

First, since the projection data only consisted of tracer injections in the right hemisphere, we flipped each of the 488 tracer images across the midsagittal plane to represent the set of projections emanating from the contralateral (left) hemisphere. Then, we computed the projection density-weighted overlap between the projection density image associated with each of the 108 seed regions considered in this experiment and the injection seed region for all 976 projection density images (488 х 2 hemispheres). The projection density-weighted overlap was computed as

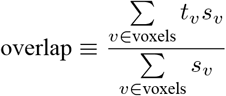

where *t_v_* is the tracer projection density value and *s_v_* is the Allen Institute defined injection fraction value at voxel *v*. For each of the 108 seed regions, the estimated polysynaptic connectivity image was constructed by choosing all projection density images corresponding to seed regions with an overlap of greater than 0.25; these images were merged voxelwise by taking the maximum projection density across overlapping images.

This image combination process was repeated to generate an estimate of polysynaptic connectivity mediated across two synapses (“2 hops”). We note that the seed regions corresponding to the complete set of 976 projection density images cover only about 30% of grey matter in the whole mouse brain, and therefore the polysynaptic connectivity images likely miss some projection tracts.

#### Transcriptomic similarity 3D datasets

Transcriptomic similarity was computed voxel-wise as the Pearson correlation coefficient between expression image voxel values and the mean expression value within the seed across all 4345 gene expression images. This resulted in 108 transcriptomic similarity images that describe the extent that voxels across the brain share similar gene expression profiles to the seed. As in the parcellation-based analysis (Section 2.5), expression images were not preprocessed in any way. Indeed, the transcriptomic similarity images computed voxelwise were spatially smooth and free of any artefacts.

#### Distance 3D datasets

For each of the 108 injection experiments considered, distances between each voxel in the brain and the boundary of the seed was computed. These distances were computed using via the fast marching method (Sethian, 1996) using Python/scikit-fmm inside a mask of the brain, emanating from the zero contour set as the boundary of the injection region.

#### Voxelwise structural covariance 3D datasets

Since the tracer connectivity data shows fine neuronal tracts, comparing these to large-scale covariance patterns determined through parcellation-based methods is not ideal. Therefore, a voxelwise approach in which structural covariance patterns are localized to specific voxels is warranted.

For each of the 108 injection sites as seed regions of interest, voxelwise structural covariance images were constructed by correlating log Jacobian determinants at each voxel in the brain with the mean of the log Jacobian determinants of voxels in the seed region. Log-transformed Jacobian determinants were used for computing correlations in order to reduce skewness in their distribution (Leow et al., 2007). As with the parcellation-based values, relative volumes were used by computing Jacobians based only on the nonlinear part of the transformations. This also avoids spurious correlations driven only by variations in whole-brain volume.

#### 3D voxelwise data comparisons and statistical methods

The 108 datasets, each corresponding to a seed region of interest, comprised of a tracer projection density image that shows monosynaptic connectivity, two polysnaptic connectivity images that estimates tracer connectivity if the tracer could hop across one and two synapses, a transcriptomic similarity image, a physical distance image, and an image of structural covariance to the seed. Structural covariance was assessed in a population of 153 mice, well above the estimated 30-40 suggested by Pagani and colleagues as necessary for reliable covariance maps (Pagani et al., 2016). Figure 2 shows the data for one of the 108 seed regions (the medial mammillary nucleus).

**Figure 2:**
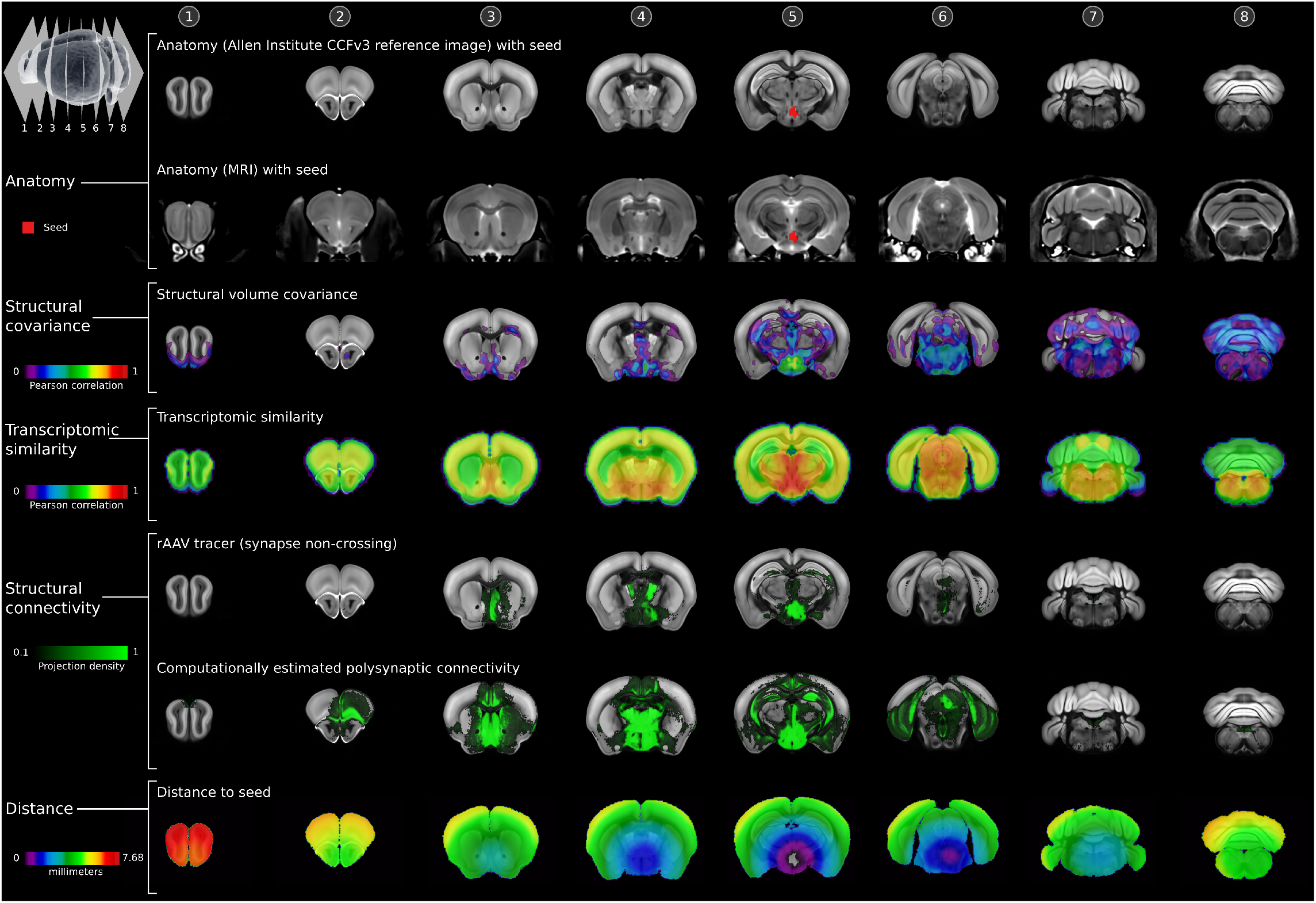
An example of the voxelwise data used to assess the relationship between structural covariance, transcriptomic similarity, connectivity, and distance. Shown are coronal slices through the mouse brain, from anterior to posterior ends, for structural volume covariance to the medial mammillary nucleus, transcriptomic similarity to that region, distance to that region, and connectivity to that region. Top two rows show anatomy (row 1: Allen Institute CCFv3 reference average template, row 2: MRI average), with seed region outlined in red. Row 3 shows structural volume covariance to the seed region, constructed by correlating voxelwise log-transformed Jacobian determinants to the seed volume (*N* = 153 images). Row 4 shows transcriptomic similarity to the seed, computed as the voxelwise correlation expression values across the brain and the mean expression value in the seed region *(N =* 4345 genes). The next two rows demonstrate structural connectivity data (projection density data from the Allen Institute); row 5 shows the path that an anterograde rAAV neuronal tracer (which does not cross synapses) takes when injected in the seed, and row 6 shows the computationally estimated image of the tracer projection if the tracer could cross a single synapse. Lastly, row 7 shows the Euclidean distance from each point in the brain to the seed.

For each of the 108 datasets, linear models were fit between structural covariance voxel values (Pearson correlations) and voxel values for monosynaptic connectivity, estimated polysynaptic connectivity (“1 hop” and “2 hops”), transcriptomic similarity (Pearson correlation), distance, and various combinations of the aforementioned predictors. Since the Allen Institute connectivity experiments consisted of injections only in the right hemisphere, these linear models were fit using voxels from the right (ipsilateral) and left (contralateral) hemispheres and compared separately. Additionally, voxels within the seed region were not considered to avoid selection bias. Each linear model was fit to approximately 2 million voxels. The coefficient of determination of each linear model, adjusted for multiple predictors (i.e. adjusted *R*^2^), was used as a measure of the variation in structural covariance values explained by the predictors. A total of 24 linear models for different combinations of predictors were fit (2 hemispheres × (5 univariate predictors + 5 bivariate predictors + 2 trivariate predictors)). Tables 2 and 3 list all the models.

**Table 2:** Variation in structural covariance, explained by univariate models. P-values reported are corrected for multiple testing as specified by Benjamini and Yekutieli (Benjamini & Yekutieli, 2001).

| Model predictors | Variation explained<br>Ipsilateral hemisphere |  | Variation explained<br>Contralateral hemisphere |  |
| --- | --- | --- | --- | --- |
|  | Bootstrapped median | Range | Bootstrapped median | Range |
| Distance | 17% ($p < 0.0001$) | 0-49% | 10% ($p = 1.0$) | 0-45% |
| Transcriptomic similarity | 13% ($p < 0.0001$) | 0-28% | 11% ($p = 0.0102$) | 0-27% |
| Monosynaptic connectivity | 12% ($p < 0.0001$) | 1-28% | 4% ($p < 0.0001$) | 0-18% |
| Polysynaptic connectivity<br>(1 hop) | 15% ($p < 0.0001$) | 0-32% | 7% ($p < 0.0001$) | 0-26% |
| Polysynaptic connectivity<br>(2 hops) | 14% ($p < 0.0001$) | 0-34% | 9% ($p < 0.0001$) | 0-36% |

**Table 3:** Variation in structural covariance, explained by multivariate models including interaction terms. In models that contain the polysynaptic connectivity term, the “1 hop” variant was used. P-values reported are corrected for multiple testing as specified by Benjamini and Yekutieli (Benjamini & Yekutieli, 2001).

|  | Variation explained<br>Ipsilateral hemisphere |  | Variation explained<br>Contralateral hemisphere |  |
| --- | --- | --- | --- | --- |
| Model predictors | Bootstrapped median | Range | Bootstrapped median | Range |
| Distance ×<br>Transcriptomic similarity | 33% (p<0.0001) | 1-60% | 26% (p<0.0001) | 1-53% |
| Distance ×<br>Monosynaptic connectivity | 25% (p<0.0001) | 3-52% | 13% (p=0.4040) | 0-48% |
| Transcriptomic similarity ×<br>Monosynaptic connectivity | 22% (p<0.0001) | 3-38% | 16% (p<0.0001) | 2-29% |
| Distance ×<br>Polysynaptic connectivity | 27% (p<0.0001) | 3-57% | 15% (p=0.0041) | 2-48% |
| Transcriptomic similarity ×<br>Polysynaptic connectivity | 24% (p<0.0001) | 4-44% | 18% (p<0.0001) | 2-34% |
| Distance ×<br>Transcriptomic similarity ×<br>Monosynaptic connectivity | 36% (p<0.0001) | 6-61% | 28% (p<0.0001) | 6-54% |
| Distance ×<br>Transcriptomic similarity ×<br>Polysynaptic connectivity | 37% (p<0.0001) | 6-63% | 29% (p<0.0001) | 6-54% |

Distributions (each comprising of 108 *R*^2^ values) representative of the variation explained by the models were tested for significance using a permutation test in which seed region labels were permuted 100000 times for each model. Additionally, the data were bootstrapped (i.e. resampled with replacement) 100000 times to generate a distribution of median values which provide intervals of confidence. The p-value for each model was assessed as the proportion of permutation-obtained medians that were greater than the 5th percentile of the bootstrapped distribution of medians. P-values were further pooled together across the 38 different models and corrected for multiple comparisons using the false discovery rate method as specified by Benjamini and Yekutieli (Benjamini & Yekutieli, 2001).

Seed regions were further clustered based on the variation in structural covariance that could be explained by distance, monosynaptic connectivity, and transcriptomic similarity. For each seed, feature vectors consisting of the three *R*^2^ values associated with the three aforementioned models were hierarchically clustered (using average linkage) into four clusters. Cluster number was determined via a scree plot. The null distribution obtained from the unclustered data (by permuting the seed region labels 100000 times for each model) was again used to calculate p-values; as before, the p-value for each model and cluster was calculated as the proportion of permutation-obtained medians that were greater than the 5th percentile of the bootstrapped distribution of medians. A total of 96 p-values (2 hemispheres × (5 univariate predictors + 5 bivariate predictors + 2 trivariate predictors) × 4 clusters) were corrected for multiple testing using the Benjamini and Yekutieli method (Benjamini & Yekutieli, 2001).

Lastly, distributions of variation explained (*R*^2^) values were examined for dependencies on tracer image properties and on variance of seed region volumes.

## 3. Results

### 3.1. Parcellation-based exploration

We first used an atlas to define structures over which a matrix of volume correlations was calculated, and compared this structural covariance matrix (Figure 3a) to similarly constructed matrices for transcriptomic similarity (Figure 3b), structural connectivity (Figure 3c), and source-target distance (Figure 3d).

**Figure 3:**
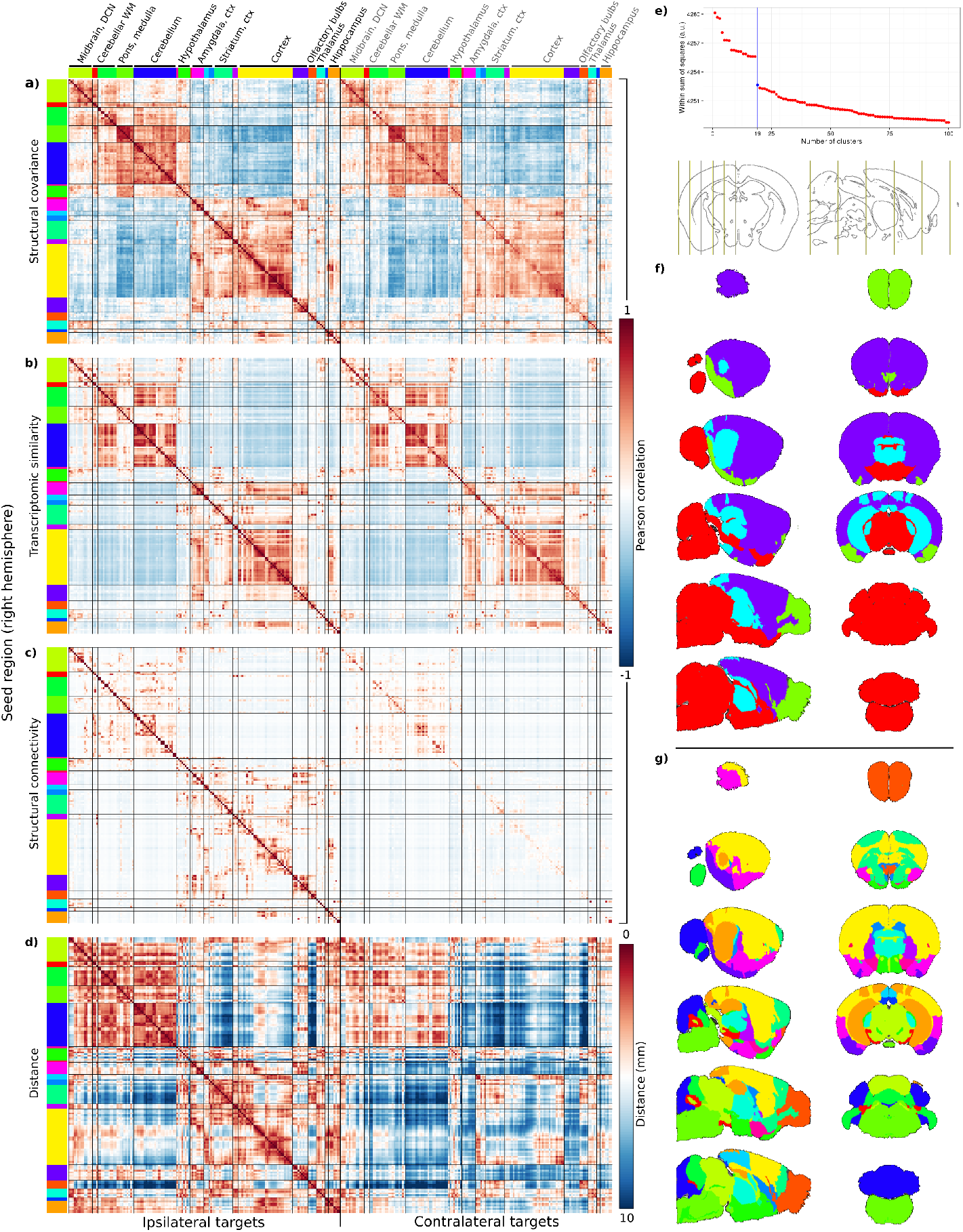
A region-based comparison of **(a)** structural covariance to **(b)** transcriptomic similarity, **(c)** structural connectivity, and **(d)** source-target distance. Rows and columns of each matrix denote atlas-defined structures, and each matrix element quantifies the association between the row-column structure pair (Pearson correlation for structural covariance, transcriptomic similarity, and structural connectivity; millimeters for distance). Rows map to source structures in the right hemisphere, while columns identify target structures ipsilateral (left half of each matrix) and contralateral (right half matrix) to the source. Structural covariance data was hierarchically clustered into 19 clusters (number of clusters determined via a scree plot **(e)**; at 19 clusters, increasing the number of clusters did not increase the within-sum-of-squares cluster distance as much as for lower number of clusters). Coarse scale clustering of structural covariance (hierarchically clustered into 4 clusters) is shown in **(f)** with arbitrary colours. All matrices were ordered according to structural covariance clustering for 19 clusters. Colour bands flanking rows and columns identify the cluster in which each region row/column lies within, the same colours identify the regions on sagittal and coronal slices of the mouse brain **(g)**. Labels at the top of the matrices indicate major structures that lie within each cluster.

#### Transcriptomic similarity, structural connectivity, and distance correlate with structural covariance

A visual inspection of matrices in Figures 3a-d indicates a correspondence between structural covariance and transcriptomic similarity, structural connectivity, and distance. At a coarse scale, strong cortex-cortex and cerebellum-cerebellum structural covariance are seen, but cortical regions generally do not correlate positively with cerebellar structures. Other notable correlations are between pons, medulla, and other nuclei nearby, including the pontine and cuneate nuclei.

A particularly strong concordance with transcriptomic similarity is seen at the whole-brain scale. For example, the structural covariance within the cerebral cortex (yellow labels) and cerebellar lobules (green and blue labels) share similar transcriptomic similarity and covariance profiles. A partial least squares decomposition and subsequent comparison of the structural covariance and transcriptomic similarity matrices with the first component results in an *R*^2^ value of approximately 50% for the structural covariance matrix, and approximately 54% for the transcriptomic similarity matrix; this component roughly outlines the separation of cortical and cerebellar structures (Figure S1). Not every pair of regions strongly correlated in volume also share similar gene expression profiles—structural covariance between hindbrain (medulla, pons) and cerebellum was not accompanied by transcriptomic similarity for example. Conversely, no pairs of structures with strong transcriptomic similarity but weak structural covariance were readily identified.

Structural covariance patterns also reflect structural connectivity organization (as described by the correlation matrix in Figure 3c), albeit to a much weaker extent. Connectivity patterns are much sparser, though clusters of structurally connected regions that also strongly covary in volume together can be readily identified. Connectivity-structural covariance concordance is stronger in the ipsilateral hemisphere. The structural connectivity matrix also resembles the transcriptomic similarity matrix in that similar clusters of regions can be visually identified, indicating that some variation in structural covariance that is explained by transcriptomic similarity could be shared by structural connectivity.

Source-target distance also correlates with structural covariance. In general, structures closer together tend to correlate more strongly in volume, although exceptions to this rule are the cuneate nucleus and medial septum, which have a correlation coefficient of ~0.5 but are relatively distant to each other, and the flocculus and paraflocculus in the cerebellum, which correlate weakly but are quite close to each other.

Figure 3 suggests that structural covariance patterns are predominantly bilateral, with the correlation structure to contralateral regions mirroring ipsilateral correlations. Although some connections are weaker (particularly contralateral cortex-cortex correlations). This bilateral covariance is reflected in the transcriptomic similarity matrix. Structural connectivity and distance matrices are also bilateral at the whole-brain scale (the two largest clusters of connected regions are preserved), but deviate at the level of individual structures, with cortical structures showing the largest bilateral differences.

#### Regions cluster into a hierarchy of neuroanatomical systems based on structural covariance patterns

We observed that hierarchical clusters of regions which covary in volume emerge. A *scree* plot (within-sum-of-squares (WSS) cluster distances plotted against cluster number) quantifies the emergence of these hierarchies as plateaus followed by drops in the WSS as the number of clusters is increased (Figure 3e). Anatomical clustering at a coarse scale (four clusters) is shown in Figure 3f. The four clusters can be labeled as: olfactory bulb and amygdalopiriform areas, cerebral cortex and striatum, hypothalamus and hindbrain, and thalamus and hippocampus.

Increasing the cluster number decreases the WSS until 19 clusters at which the WSS plateaus; the four matrices were thus split and ordered into 19 clusters, with regions lying in the same cluster being grouped together. Row and column colour bands flanking the correlation matrices represent the cluster to which each region is assigned; the same colours are used to show this clustering in anatomical space in Figure 3g. At this finer scale, clusters formed are contiguous; regions most strongly coupled together in their volume are also neighbouring regions. Clusters vary in size (both in the number of regions contained and in volume of the brain covered), ranging from the large cortical cluster of similar covariance patterns (yellow) to the single region cluster consisting of the basal forebrain (pink).

### 3.2. Seed-based voxelwise analysis

In the seed-based analysis, the associations between structural covariance and transcriptomic similarity, structural connectivity, and distance to seed, were assessed voxel-wise for each seed by fitting linear models.

#### Variation explained by univariate models

Table 2 shows the variation explained by distance, transcriptomic similarity, and connectivity, across the 108 large seeds chosen and over voxels in the hemisphere ipsilateral to the seed regions (right hemisphere). In general, variation explained in the ipsilateral hemisphere was slightly higher than in the contralateral hemisphere. The median variation explained values under all models were highly unlikely to be explained by chance (p<0.0001 for all models, except p=0.00033 for distance in the contralateral hemisphere). Since the “2 hop” estimated polysynaptic connectivity predictor did not explain much more variation than its “1 hop” counterpart in either hemisphere, it was not used in any further analyses.

#### Variation explained by multivariate models

Overlap in the explanatory value of the predictors was assessed through multivariate linear models that included interactions. Table 3 shows the variation explained by combinations of the predictors, across the 108 large seeds chosen and over voxels in separate hemispheres. Bivariate models explain more variation than univariate models. Apart from the predictors explaining slightly less variation, particularly for models involving distance, trends in the contralateral hemisphere mirror that of the ipsilateral hemisphere. Figure 4 shows the variation explained by all models (univariate and multivariate).

**Figure 4:**
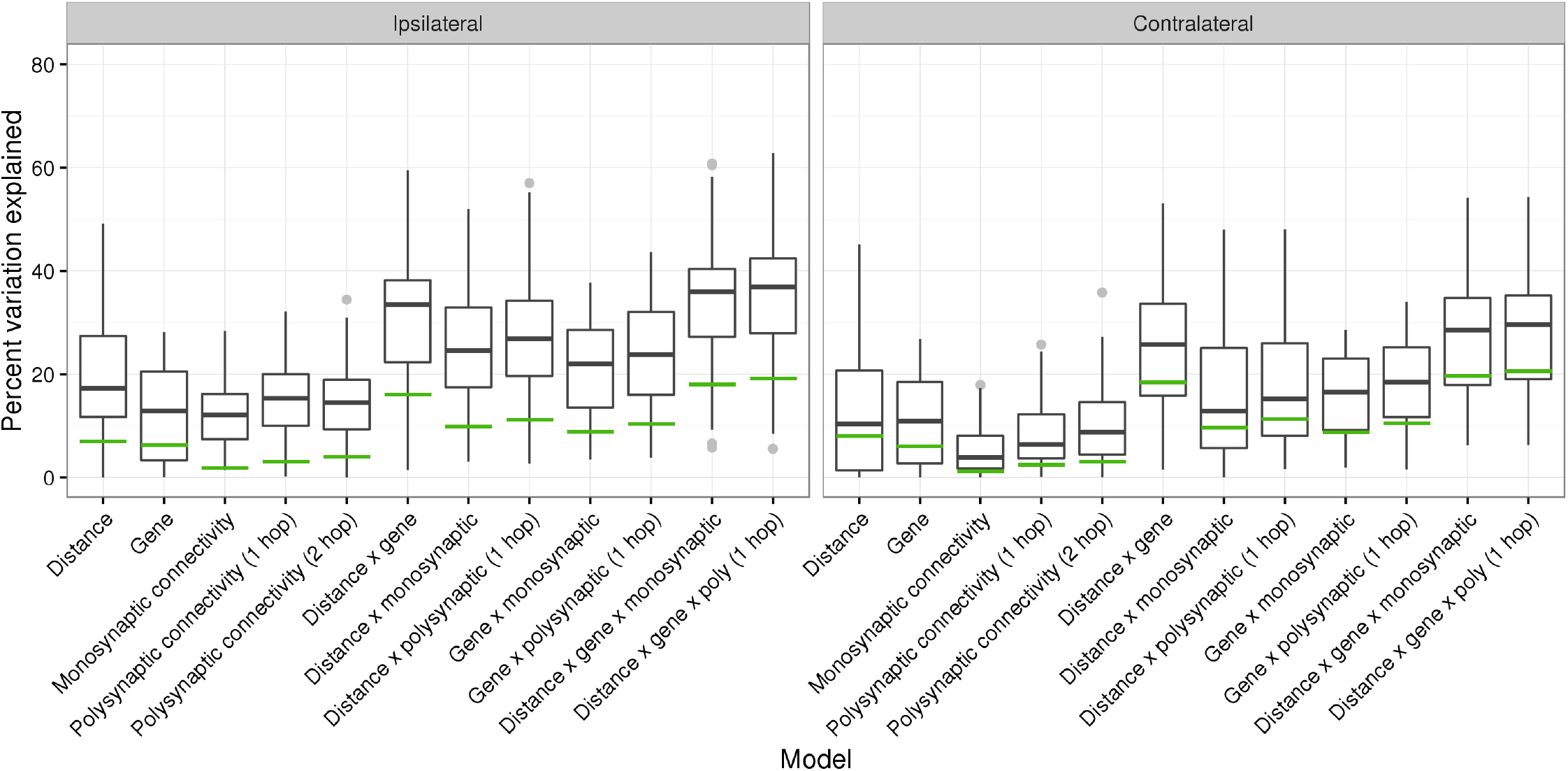
Variation in structural covariance explained by distance to seed, transcriptomic similarity to seed, and connectivity (monosynaptic and estimated polysynaptic) to seed, for 108 seed regions. Variation explained (*R*^2^) was computed separately over voxels in the right (ipsilateral) and left (contralateral) hemispheres. A permutation test was performed to determine the distribution of medians under the null hypothesis; the green line is the value above which 5% of medians lie under the null hypothesis.

#### Variation explained by connectivity does not depend on tracer properties

To ensure that explained variation values are not due to tracer experiment confounds, we examined whether explained variation for each seed correlated with tracer volume, maximum distance the tracer projects, estimated polysynaptic tracer volume, and the maximum polysynaptic tracer distance (Figure S3a,b,d,e). We found that the variation explained by monosynaptic connectivity did not depend on tracer volumes or the maximum distances that the tracers projected; a similar lack of relationship held for polysynaptic connectivity. We also verified that injection volume did not affect variation explained by monosynaptic connectivity, thus validating our seed choice criteria (Figure S3c).

#### Variation explained by expression similarity is correlated with transcriptomic commonness

Lastly, we defined the transcriptomic commonness of a seed region as the sum of transcriptomic similarity correlation coefficients over all voxels in the brain, multiplied by the voxel volume. Noting that this measure represents the uniqueness of transcriptome (the higher the transcriptomic commonness, the less spatially unique the transcriptomic similarity pattern is) and is not necessarily a confound, we found that the variation explained by transcriptomic similarity depends on transcriptomic commonness (Figure S3f). Given that cortical and subcortical regions share different gene expression and explained variation patterns, we examined cortical and subcortical seeds separately and found that variation explained by transcriptomic similarity correlates more strongly with transcriptomic commonness in the cortex than subcortex.

#### Seed regions cluster into distinct neuroanatomical systems based on patterns of explained variation

To examine whether structural covariance is explained by transcriptomic similarity, connectivity, and distance differently based on location of the seed, we clustered the seed regions into four groups via hierarchical clustering, using the variation explained by distance, transcriptomic similarity, and monosynaptic connectivity to each seed as a three dimensional vector associated with each seed (Figure 5). The four clusters consist of seeds distributed in a spatially unique patterns, and map to unique explained variation trends. These are as follows,

- Cluster A (43 seeds located primarily in the midbrain, posterior cortex/visual areas, and posterior hypothalamus): distance, transcriptomic similarity, and connectivity each explain equal amounts of variation (~14-18%, more than chance) in the ipsilateral hemisphere.
- Cluster B (22 seeds, located primarily in the anterior and posterior hypothalamus): distance and transcriptomic similarity explain almost no variation (<5%), connectivity explains some variation (~3-8%) above chance in the ipsilateral hemisphere.
- Cluster C (23 seeds, located primarily in the hindbrain): transcriptomic similarity explains almost no variation, most variation is explained by distance (~25%) although connectivity also has a role (~12-17%). Distance and connectivity explain structural covariance in the ipsilateral hemisphere more so than can be explained by chance alone.
- Cluster D (20 seeds, located primarily in the anterior cortex): distance explains the most variation by far (~40%), but transcriptomic similarity and connectivity also explain structural covariance more than chance can (~9-22%) in the ipsilateral hemisphere.

**Figure 5:**
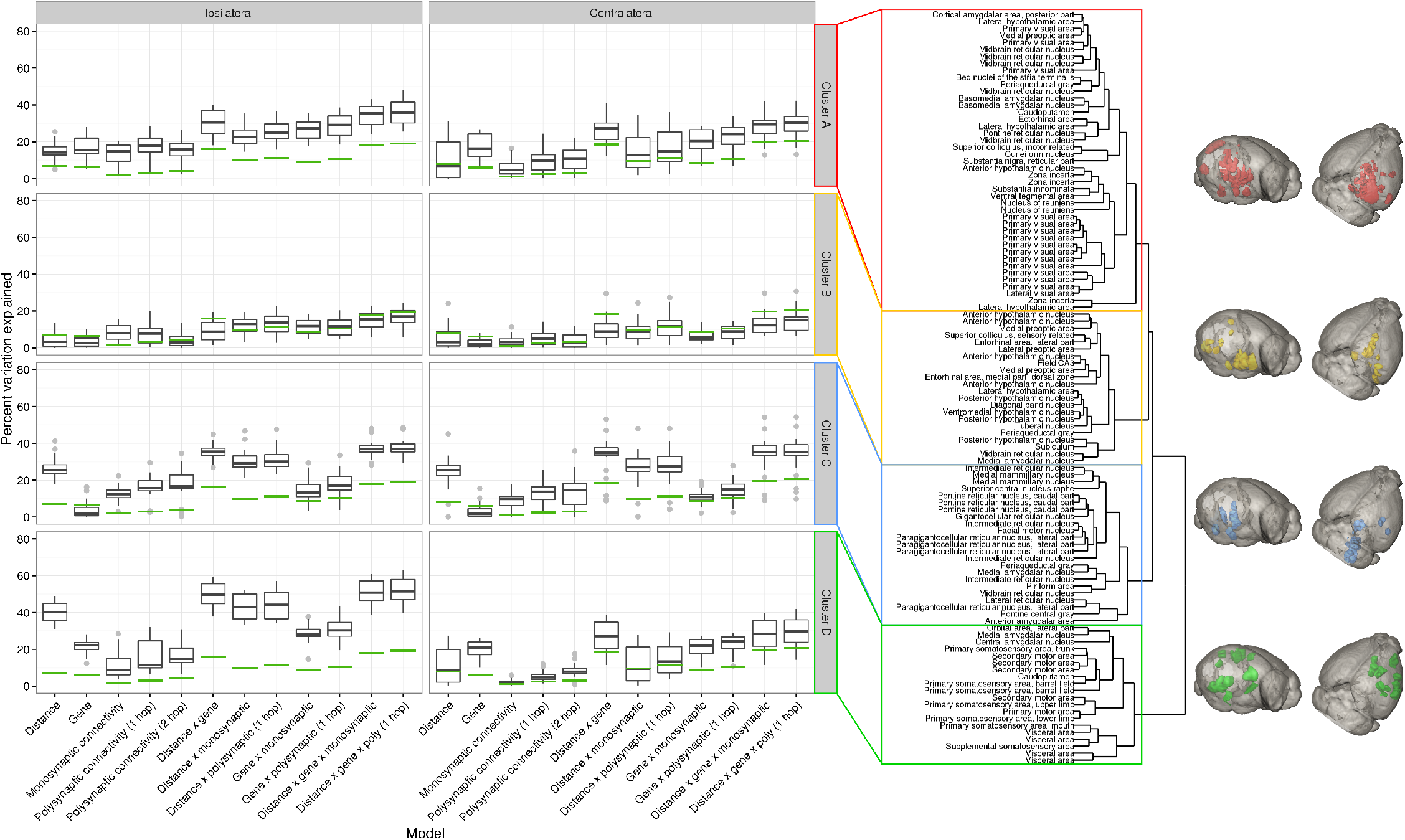
Variation explained for 108 seed regions, split into four clusters based by hierarchically clustering *R*^2^ values in the ipsilateral hemisphere for distance, transcriptomic similarity, and monosynaptic connectivity. The dendrogram corresponding to the clustering is shown to the right of the boxplots, and the spatial location of seeds within each cluster are rendered on the right.

These results suggest that transcriptomic similarity is primarily associated with structural covariance to the cortex, whereas variation explained by connectivity is less localized, and is particularly high for hindbrain regions.

#### Variation explained does not depend on the variance in volumes of seed regions

We examined whether the lack of variation explained for seed regions in Cluster B could be attributed to low variance in volumes of those seed regions. If a certain amount of variance in seed region volumes might be attributed to noise, then constructing structural covariance maps for the seeds with variance below the noise threshold will result in noise driven correlations. Variance in seed region volumes are indeed lower for seeds in Cluster B (Figure S4a), but a further investigation shows no positive correlation between variation explained values and variance in seed region volumes within clusters (Figure S4b).

#### Variation explained by distance, transcriptomic similarity, and structural connectivity demonstrate spatially nonuniform and distinct patterns

To examine brainwide patterns of explained variation, we repeated this voxelwise comparison of structural covariance to distance, transcriptomic similarity, and connectivity using every voxel as a seed, albeit at a 4x lower resolution so that computations were feasible. Not every voxel was in a seed region, we therefore used correlations over tracer images (as in the atlas-based analysis) as a measure of structural connectivity. Figure 6 shows the extent that distance, transcriptomic similarity, monosynaptic connectivity, and all three combined predictors explain structural covariance to each voxel. Broadly, transcriptomic similarity seems to best explain structural covariance to the cortex and striatum. Connectivity explains structural covariance to the cortex, striatum, and hindbrain. Distance explains most variation in frontal areas of the cortex and hindbrain; together, the three predictors explain most of the variation in cortex (cingulate, motor, somatosensory, orbital, and frontal association areas) and hindbrain (pons, medulla, and parts of the cerebellum, medial septum), and least variation in the thalamus, hypothalamus, and hippocampi.

**Figure 6:**
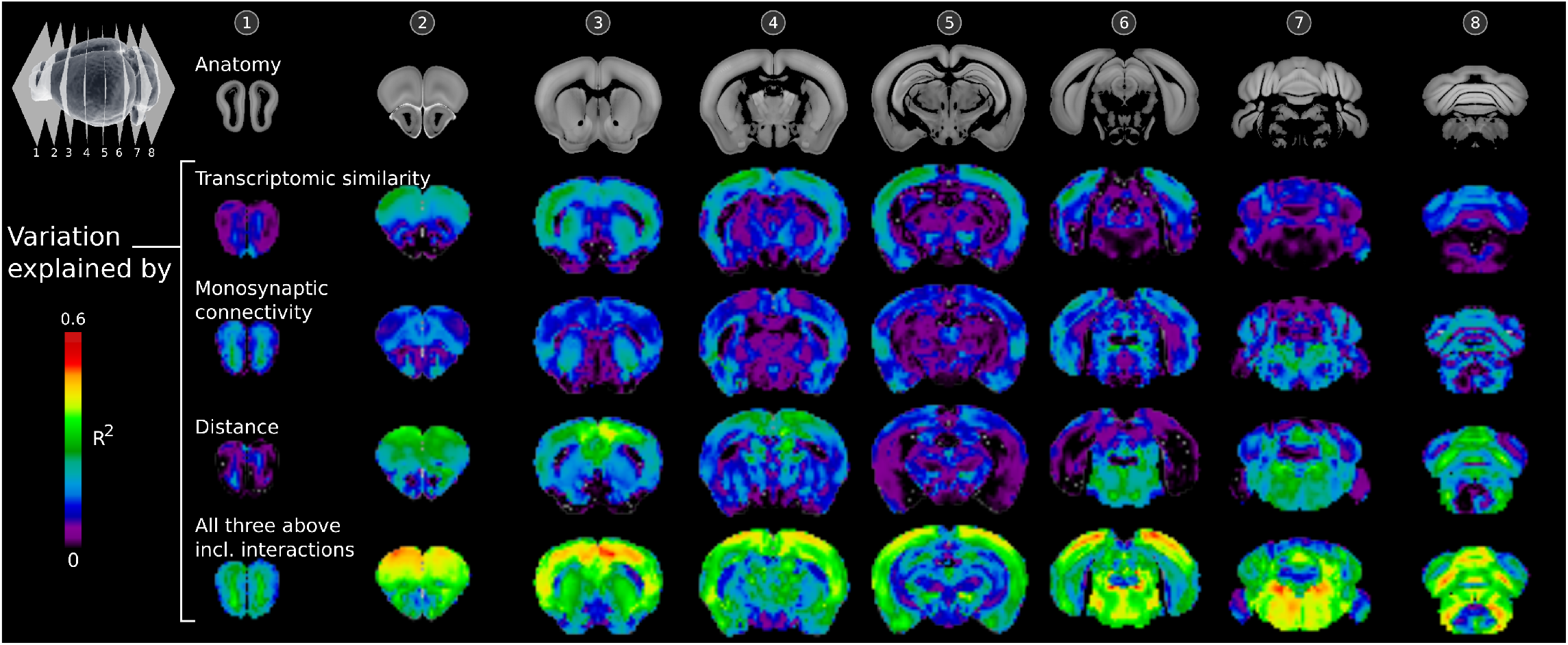
Brainwide patterns of explained variation. Shown are coronal slices through the mouse brain, with the first row displaying anatomy. The next four rows show variation explained values for structural covariance to each seed voxel plotted as a colour at that voxel. Rows correspond to variation explained by transcriptomic similarity (row 2), monosynaptic connectivity (row 3), distance (row 4), and all three previous predictors together (row 5).

Seed region data, variation explained values for all 24 models (12 × 2 hemispheres), and cluster assignment data are provided for each of the 108 seed regions in the Supplementary Table S1.

## 4. Discussion

Connectivity related plasticity and coordinated neurodevelopment (guided by spatially and temporally coordinated patterns of gene expression) are two interacting mechanisms that are thought to underlie structural covariance (Evans, 2013). Our objective was to examine the association between structural volume covariance and structural connectivity, transcriptomic similarity, and distance, and thereby provide insights into why regions couple together in their volumes.

### Comparisons to transcriptomic similarity, structural connectivity, and distance

The parcellation-based exploration shows a strong correspondence between the structural covariance matrix and transcriptomic similarity matrix, suggesting a role for transcriptomic similarity in structural covariance. Clusters of highly correlated regions within the cortex, cerebellum, and hindbrain (correlated in transcriptomic similarity and volume) connect regions of common developmental origins, pointing to the idea that the structural covariance network seen might arise from coordinated gene expression during neurodevelopment. An interesting feature of the cortex is that regions within the cortex cluster more strongly together than other pairs of regions. In the atlas-based clustering into 19 clusters, most of the cerebral cortex remained in one cluster, indicating that cortical volumes might arise from common underlying factors that spans the cortex. Recent work by Romero Garcia et al. (2017) suggests that human supragranular enriched genes might be one such factor. Longitudinal volume data along with expression data at earlier timepoints would help further probe the temporal development of structural covariance networks and determine whether structural covariance arises from coordinated expression of developmental cues during brain growth.

Structural covariance also reflects structural connectivity patterns, although this association is not as strong as with transcriptomic similarity. This might be due to the sparseness of tracers, i.e. not enough tracer experiments were considered in building a whole-brain connectivity matrix (the seeds selected covered 18% of grey matter in the right hemisphere). Nonetheless, the patterns of structural connections (mediated by projection tracts that do not cross synapses) also reflect structural covariance more than chance can explain alone. We note that connectivity and transcriptomic similarity are not necessarily mutually independent. Spatial and temporal gene expression patterns guide the development of the brain, including the formation of the structural connectome via, for example, the expression of neuron growth factors and axon guidance molecules (Plachez & Richards, 2005). Indeed, rodent connectivity can be predicted from the spatial coexpression patterns of a set of genes related to neurodevelopment (French & Pavlidis, 2011), and in the case of the cortex, age-related changes in structural covariance during adolescence are predominant in the frontal lobe, consistent with the tuning of frontal lobe structural connections during this developmental period (Vasa et al., 2017). Given that the human supragranular genes implicated in structural covariance (Romero Garcia et al., 2017) are associated with cortico-cortical connectivity (Krienen et al., 2016), structural connectivity driven by the coexpression of neuron-related genes between regions is a candidate mechanism for the coupling of volumes between those regions.

Related to structural connectivity, another measure to examine would be functional connectivity. In both humans and mice, networks of functional connections are associated with both structural connectivity (Honey et al., 2009; Grandjean et al., 2017; Mills et al., 2017) and transcriptomic similarity (Richiardi et al., 2015; Vértes et al., 2016; Mills et al., 2017). Furthermore, specific functional tasks have been shown to correlate with volumes of regions subserving those tasks in both mice (Lerch et al., 2011b) and humans (Maguire et al., 2000). Thus, functional connectivity is also expected to associate with structural covariance. Whether functional connectivity explains any more variation in structural covariance beyond the variance explained by distance, transcriptomic similarity, and distance remains to be seen.

The association between structural covariance and distance between regions is also apparent, but this link is not entirely clear. Our results show that if a region grows in volume, it does not push against and thereby compress neighbouring regions. Instead, neighbouring regions also tend to grow. This preference for structural covariance (positive correlations) at short distances might arise from the fact that nearby regions tend to share the same gene expression profiles due to their common embryonic origins, although the tendency for nearby regions to connect together (Scannell et al., 1995) might also explain high structural covariance. In constructing structural covariance maps, the registration procedure includes a regularization term which smooths the deformation fields from which the Jacobian determinants are computed. This spatial smoothing would also explain positive correlations between voxels that are very close to each other.

The voxelwise analysis quantified the link between structural covariance and transcriptomic similarity, structural connectivity, and distance by quantifying the variation in structural covariance that could be explained by the aforementioned data. Multivariate models consisting of multiple predictors tend to explain more variation than single predictors, suggesting that the explanatory values of transcriptomic similarity, connectivity, and distance add to some extent, rather than completely overlap. In the voxelwise analysis, we also examined structural connectivity mediated by synapse-separated tracts by computationally estimating what the rAAV tracer would look like if it could cross synapses. This was motivated by the observation of bilateral patterns of structural covariance, and generally weaker monosynaptic projections from seeds to contralateral areas as compared to ipsilateral areas. Considering connectivity mediated by multiple tracts connecting across synapses would also better reflect functional connectivity and explain contralateral coactivations and structural covariance. Unsurprisingly, polysynaptic connectivity explains slightly more variation than monosynaptic connectivity in the contralateral hemisphere. Interestingly, polysynaptic connectivity “hopping” across 2 synapses did not explain much more variation than the single hop variant, likely due to more of the brain being filled by the (computationally estimated) tracer, including in areas of low structural covariance.

Lastly, variation explained by structural connectivity did not depend on tracer confounds. Structural covariance to seeds with high transcriptomic commonness were explained more by transcriptomic similarity however, especially for cortical seeds, suggesting that a common set of cortical development factors might underlie covariance.

In this study, we did not address negative correlations. Negative correlations are generally weak (especially in the voxelwise images). Similar to negative correlations that arise in fMRI data when removing the global signal (Murphy et al., 2009), negative correlations seen in this study can frequently be attributed to normalization by overall brain volume.

### Spatial patterns of explained variation

Clustering into four groups the explained variation data across 108 seeds results in distinct trends of variation explained, and these trends split seed regions into spatially distinct areas. It is important to note that the source of transcriptomic similarity data (*in-situ* hybridization) and connectivity data (projection density derived from two-photon fluorescence signal) are quite different, and this constrains comparisons on the extent that one predictor drives structural covariance in relation to the other. We can examine the variation explained by individual models across seeds, clusters, or space however. For the four clusters of seeds, transcriptomic similarity tends to explain clusters with seeds in the cortex better than others, again pointing to a role for coordinated neurodevelopment in cortical structural covariance. Which genes are involved in structural covariance, particularly in the cortex, have yet to be identified. Connectivity on the other hand plays a role in explaining structural covariance in all clusters, although explained variation is low in the hypothalamus (yellow) cluster. Interestingly, transcriptomic similarity or distance also does not play a large role in hypothalamic structural covariance. While the variances in the volumes of seed regions in the hypothalamic cluster were low, this does not explain the low variation in structural covariance explained by all models. Overall, structural covariance in the two clusters (red and green) corresponding to cortical seeds are explained to the same extent by transcriptomic similarity and connectivity, though distance has a larger role for seeds in the anterior cortex (green cluster). This suggests that the association of structural covariance to distance might not entirely be due to similar transcriptomic similarity nearby, or short range projection tracts.

The brainwide maps of explained variation largely mirror variation explained patterns seen from clustering the seeds: transcriptomic similarity is associated with structural covariance in the cortex, while connectivity is associated with structural covariance across the brain, and particularly strongly in the cortex, striatum, and hindbrain. Figure 4 of Lein et al. (2007) demonstrates that for the top 100 genes expressed in a chosen structure, the hippocampus, olfactory bulbs, cortex, and thalamus exhibit highly enriched gene expression, while the hypothalamus, midbrain, pons, and medulla exhibit spatially overlapping patterns of expression. This spatial separation of structures by their expression patterns seems to mirror the pattern of variation explained by transcriptomic similarity, suggesting that structural covariance that is linked to transcriptomic similarity might arise from a smaller set of locally enriched genes. Within specific structures, differences in explained variation might map to functional differences; for example, differences in explained variation in the dorsal and ventral striatum might reflect the different connectivity profiles (Hintiryan et al., 2016) and functions (Koenigs & Grafman, 2009) of these areas. Similarly, structural covariance to different nuclei in the thalamus are explained to different extents by transcriptomic similarity and connectivity. These results were unexpected; we had hypothesized that connectivity would explain structural covariance better in the cortex (typically considered to be more plastic than hindbrain structures), while transcriptomic similarity would explain structural covariance better in the less-plastic and developmentally older subcortical and hindbrain regions.

### What explains the rest of the variation?

Even if structural covariance was perfectly correlated with transcriptomic similarity or structural connectivity, noise introduced by data acquisition and processing would result in an imperfect correlation. For instance, registration of mouse MR images does not perfectly recover volume differences, particularly for small or non-compact structures (van Eede et al., 2013). Potential explanations for this missing variation beyond noise could be both data related (i.e., the data does not capture all sources of variation) and model related (linear models might underfit the data). A data-related constraint was that we used gene expression data which quantified expression levels at around postnatal day 60 of the mouse, while critical periods of brain development are notably missed. Given that coordinated neurodevelopment through these early timepoints shape the volumes used to construct structural covariance maps in this cross-sectional study, we suspect that if a similar analysis was performed with gene data through development, transcriptomic similarity might explain a larger amount of variation in structural covariance. As for the latter point, underestimating explained variation might arise from the use of linear—rather than non-linear—models. Our model assumes a linear response of structural covariance to transcriptomic similarity, connectivity, and distance. It is not difficult to imagine that transcriptomic similarity or connectivity might induce a more discrete transition in structural covariance; for example, below a certain transcriptomic similarity threshold, similarity in the expression profiles might not result in structural covariance and vice versa. Although the data might be underfitted by our assumption of linearity, we chose to use linear models because of the simple interpretation of the coefficient of determination *R*^2^ (adjusted for multiple predictors) as variation explained. Analogues of the *R*^2^ value exist for non-linear models (e.g. the McFadden *R*^2^ (McFadden, 1974)), but are thought to underestimate variation explained (Domencich & McFadden, 1975). Lastly, we note that structural covariance was computed in a group of inbred C57Bl/6 mice. We hypothesize that in outbred strains, increased genetic heterogeneity might induce stronger transcriptomic similarity-associated structural covariance between regions.

### Conclusions and future considerations

In this study, we show that structural covariance is explained by transcriptomic similarity, structural connectivity, and distance more so than chance alone. Given the neuronal tracer data as a representation of structural connectivity underlying plasticity (regions that “fire together, wire together, grow together”) and transcriptomic similarity images as a model of coordinated neurodevelopment, our results suggest a role for both connectivity driven plasticity and coordinated neurodevelopment in the coupling of structures in their volumes. The extent to which these mechanisms drive structural covariance varies across the brain however, with cortical and subcortical structures showing different patterns of variation explained by structural connectivity, transcriptomic similarity, and distance. Our results support previous findings that structural covariance patterns closely mirror patterns of coordinated neurodevelopment, and that covariance is related to (but is not fully explained by) structural connectivity. Together with the aforementioned studies, these results point to a role for structural covariance in the search for biomarkers of disease and treatment response in neurodevelopmental or connectivity disorders such as autism. The exploratory analysis that we carried out might help focus future biomarker searches to specific regions of the brain—structural covariance studies on disorders of gene expression might be better suited in examining cortical volumes, although if aberrant connectivity is involved, other brain areas such as the hindbrain might also be of interest.

## Footnotes

1 at the time this study was conducted

## Acknowledgments

We thank the Ontario Brain Institute (OBI), Canadian Institutes of Health Research (CIHR), and Restracomp (SickKids Research Training Centre) for funding support. We also thank the Allen Institute for Brain Science for providing connectivity (©2011 Allen Institute for Brain Science. Allen Mouse Brain Connectivity Atlas. Available from: connectivity.brain-map.org) and gene expression (©2004 Allen Institute for Brain Science. Allen Mouse Brain Atlas. Available from: mouse.brain-map.org) data used in this study. Computations were performed on the gpc supercomputer at the SciNet HPC Consortium. SciNet is funded by: the Canada Foundation for Innovation under the auspices of Compute Canada; the Government of Ontario; Ontario Research Fund - Research Excellence; and the University of Toronto.

---

**Figure S1:**
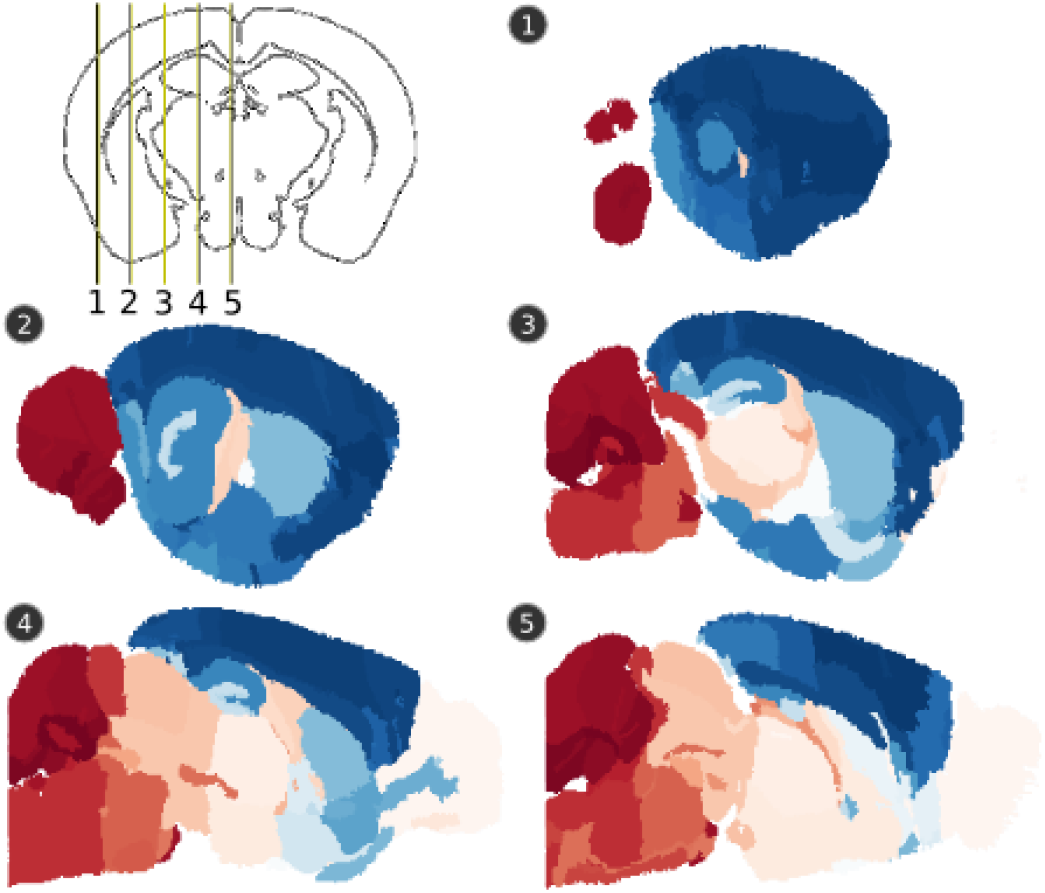
A visualization of the first component obtained from a partial least-squares decomposition of transcriptomic similarity and structural covariance matrices, plotted on sagittal slices of the mouse brain. Regions are coloured (red: positive, blue: negative) according to the value of the corresponding element in the component. These images show a separation of the cortex and anterior areas of the brain from the hindbrain and posterior areas that is common to both structural covariance and transcriptomic similarity matrices.

**Figure S2:**
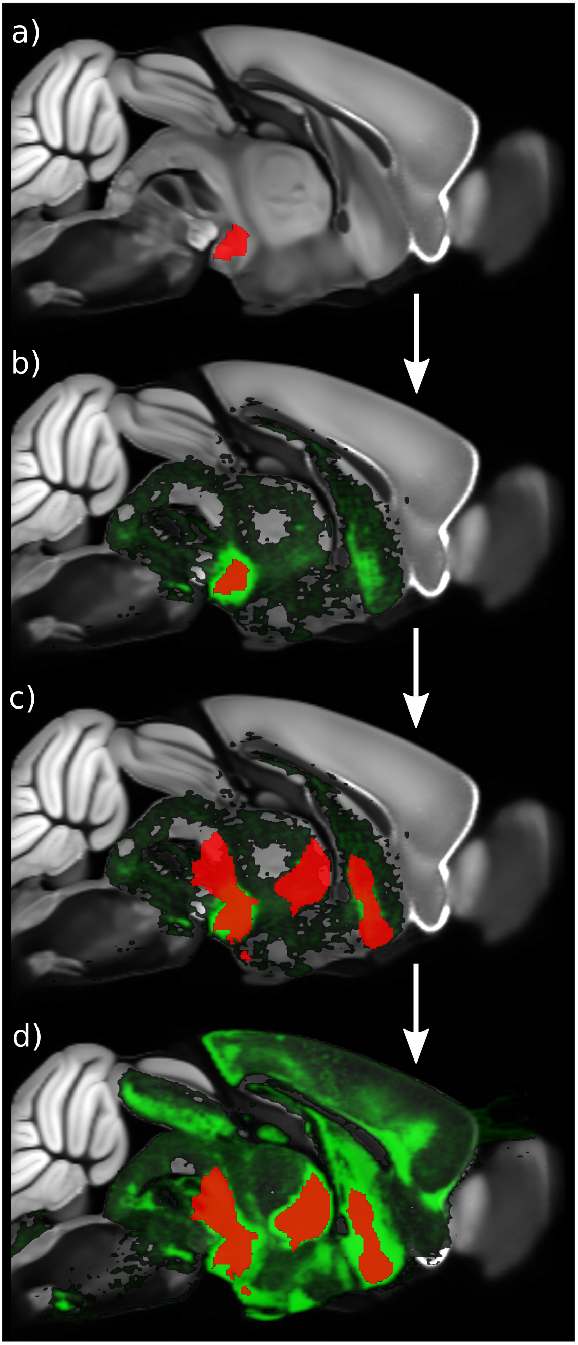
An illustration of the method used to estimate where the tracer would project if it were allowed to cross a single synapse. Shown are sagittal slices of the mouse brain close to the midline, demonstrating: (a) the original injection region, (b) the original tracer which does not cross synapses, (c) other injection regions which overlap the original tracer (by at least 25% of overlapping injection regions’ volume), and (d) the original tracer merged with tracer projections emanating from the overlapping injection sites (merging was performed voxelwise as the maximum value across tracer projection density images).

**Figure S3:**
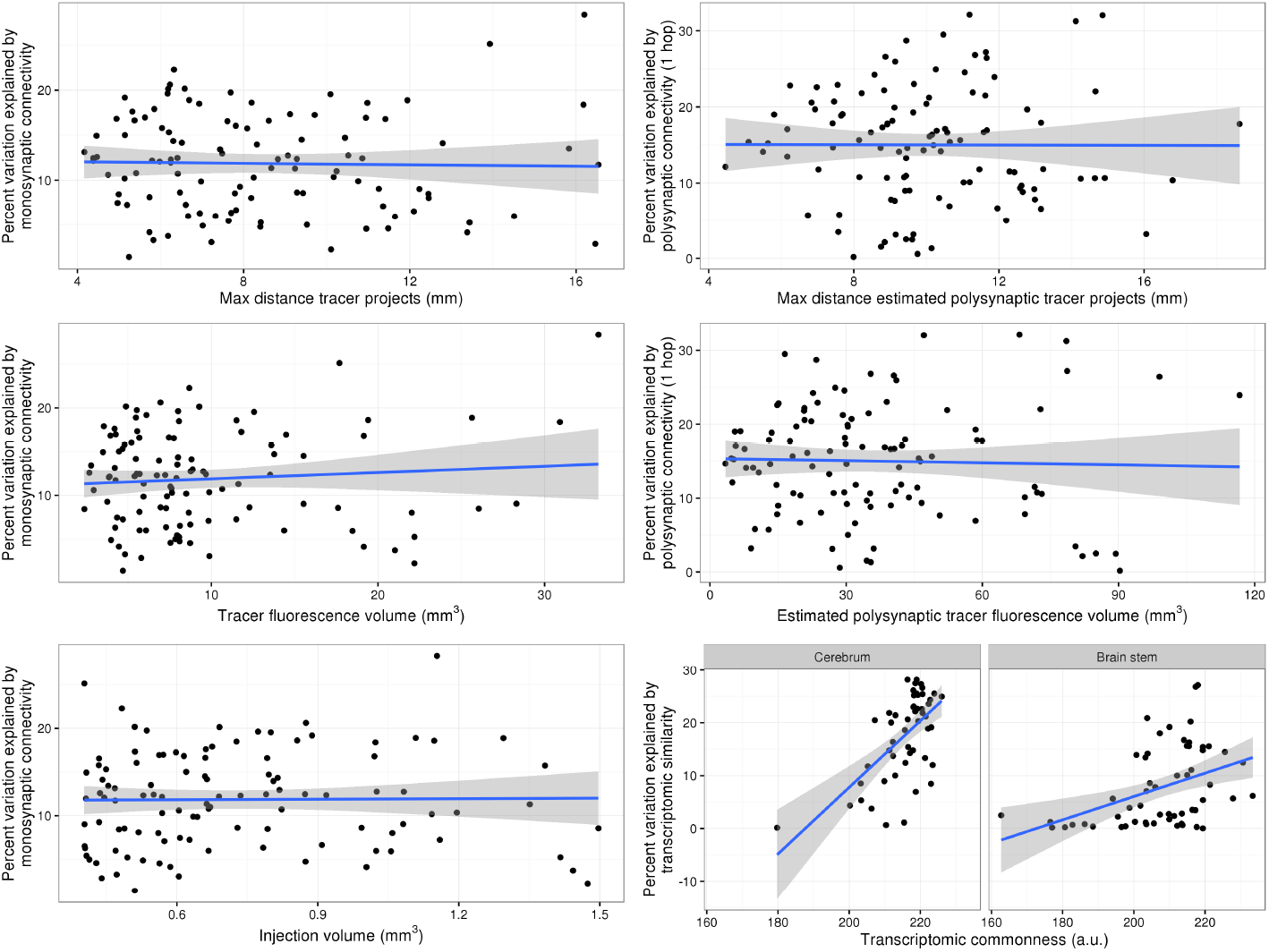
Dependence of explained variation (R^2^) on tracer injection and expression image properties. Shown (for the 108 seed regions) are (a) variation explained by monosynaptic connectivity as a function of maximum tracer projection distance, (b) variation explained by polysynaptic connectivity as a function of maximum polysynaptic projection distance, (c) variation explained by monosynaptic connectivity as a function of tracer fluorescence volume, (d) variation explained by polysynaptic connectivity as a function of polysynaptic tracer volume, (e) variation explained by monosynaptic connectivity as a function of injection volume, and (f) variation explained by transcriptomic similarity as a function of transcriptomic commonness, split by seed location in the cerebral cortex [Allen Institute classification: “cerebrum”] and subcortex/hindbrain [Allen Institute classification: “brain stem”]. Transcriptomic commonness is a measure of the uniqueness of the gene expression profile of the seed measured across voxels, i.e. the higher the transcriptomic commonness, the more voxels share a similar expression profile as the seed, and therefore the less unique the expression profile is within the seed.

**Figure S4:**
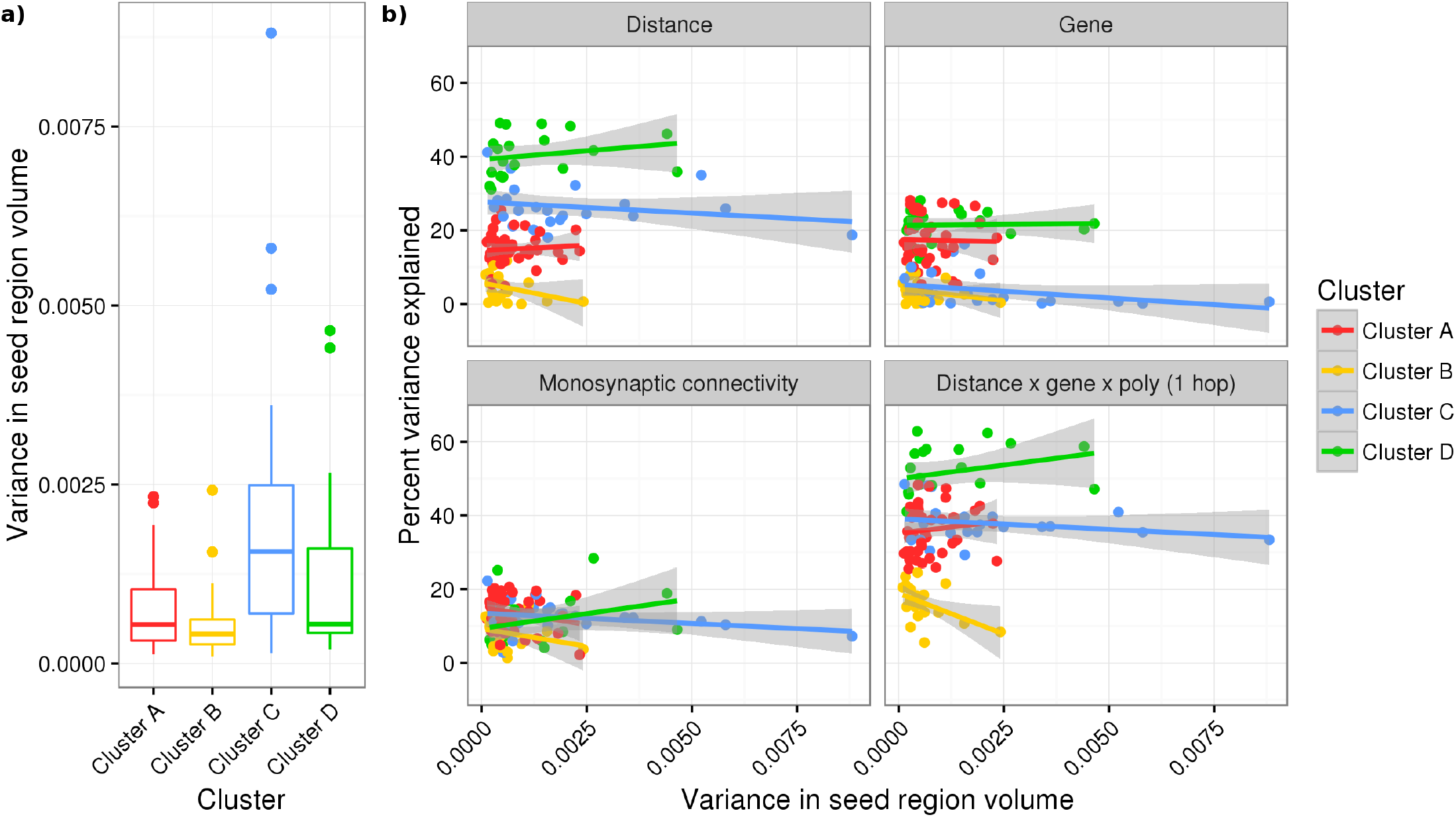
Dependence of explained variation *(R*^2^) on the variance in seed region volume. In **(a)**, the distribution in variance across seeds are plotted for all 108 seeds split into 4 clusters; seeds in Cluster C (hindbrain) vary the most in volume whereas seeds in Cluster B (hypothalamus) vary the least. In (b), the variation explained by four models (distance, transcriptomic similarity, monosynaptic connectivity, and the full model of all different types of predictors) is plotted against this variance in seed region volume for 108 seeds, split across four panels corresponding to the models. Additionally, datapoints are coloured to reflect the cluster in which each seed region lies.

Table S1: Information on the 108 seed regions, along with variation explained by each of the 24 models (12 х 2 hemispheres) for each seed.

## Supplementary Data

Processed data described in this manuscript are contained in a .RData file, `manuscript data.RData`, attached as a gzipped archive in the supplementary materials. The data can be loaded in R via load(). The data is provided in a nested list structure, and includes

- Processed data used in the parcellation analysis
  - volumetric data
    * brain region volumes, for 153 mice
    * gene expression in each region, for 4345 expression images
    * tracer projection density mean value in each region, for 976 connectivity datasets (488 experiments with seed region in each of the two hemispheres)
    * estimated polysynaptic projection density mean value in each region, for 976 connectivity datasets (488 experiments with seed region in each of the two hemispheres)
  - matrix data
    * structural covariance (correlation) matrix
    * transcriptomic similarity matrix
    * monosynaptic connectivity matrix
    * polysynaptic connectivity matrix
    * distance matrix, for Euclidean distance between centroids of regions
  - atlas structure names and cluster assignments
  - Allen Institute-associated experiment IDs for 488 connectivity experiments considered
  - Gene names and Allen Institute-associated experiment IDs for the 4345 coronal expression datasets used

- Processed data used in the seed-based voxelwise analysis
  - seed region information and associated variation explained values for each of the 24 models (same as Supplementary Table S1)
  - summaries of variation explained values for each of the 24 models, including bootstrapped estimates of the median variation explained and p-values

## References

Alexander-Bloch, A., Giedd, J. N., & Bullmore, E. (2013a). Imaging structural co-variance between human brain regions. Nature Reviews Neuroscience, 14, 322–336.

Alexander-Bloch, A., Raznahan, A., Bullmore, E., & Giedd, J. (2013b). The convergence of maturational change and structural covariance in human cortical networks. The Journal of Neuroscience, 33, 2889–2899.

Alexander-Bloch, A. F., Reiss, P. T., Rapoport, J., McAdams, H., Giedd, J. N., Bullmore, E. T., & Gogtay, N. (2014). Abnormal cortical growth in schizophrenia targets normative modules of synchronized development. Biological Psychiatry, 76, 438–446.

Avants, B. B., Tustison, N., & Song, G. (2009). Advanced normalization tools (ANTS). Insight Journal, 2, 1–35.

Behrens, T., Berg, H. J., Jbabdi, S., Rushworth, M., & Woolrich, M. (2007). Probabilistic diffusion tractography with multiple fibre orientations: What can we gain? NeuroImage, 34, 144–155.

Benjamini, Y., & Yekutieli, D. (2001). The control of the false discovery rate in multiple testing under dependency. Annals of Statistics, 29, 1165–1188.

Bernhardt, B. C., Bernasconi, N., Hong, S.-J., Dery, S., & Bernasconi, A. (2016). Subregional mesiotemporal network topology is altered in temporal lobe epilepsy. Cerebral Cortex, 26, 3237–3248.

Bernhardt, B. C., Chen, Z., He, Y., Evans, A. C., & Bernasconi, N. (2011). Graph-theoretical analysis reveals disrupted small-world organization of cortical thickness correlation networks in temporal lobe epilepsy. Cerebral Cortex, 21, 2147–2157.

Bernhardt, B. C., Valk, S. L., Silani, G., Bird, G., Frith, U., & Singer, T. (2014). Selective disruption of sociocognitive structural brain networks in autism and alexithymia. Cerebral Cortex, 24, 3258–3267.

Bethlehem, R. A., Romero-Garcia, R., Mak, E., Bullmore, E., & Baron-Cohen, S. (2017). Structural covariance networks in children with autism or ADHD. Cerebral Cortex, 27, 4267–4276.

Bruno, J. L., Hosseini, S. H., Saggar, M., Quintin, E.-M., Raman, M. M., & Reiss, A. L. (2016). Altered brain network segregation in Fragile х Syndrome revealed by structural connectomics. Cerebral Cortex, 27, 2249–2259.

Cahill, L. S., Steadman, P. E., Jones, C. E., Laliberté, C. L., Dazai, J., Lerch, J. P., Stefanovic, B., & Sled, J. G. (2015). MRI-detectable changes in mouse brain structure induced by voluntary exercise. NeuroImage, 113, 175–183.

Docherty, A. R., Sawyers, C. K., Panizzon, M. S., Neale, M. C., Eyler, L. T., Fennema-Notestine, C., Franz, C. E., Chen, C.-H., McEvoy, L. K., Verhulst, B. et al. (2015). Genetic network properties of the human cortex based on regional thickness and surface area measures. Frontiers in Human Neuroscience, 9, 440.

Domencich, T., & McFadden, D. (1975). Statistical estimation of choice probability functions. In Urban Travel Demand: A Behavioral Analysis chapter 5. (pp. 101–25). Amsterdam: North-Holland.

Dorr, A. E., Lerch, J. P., Spring, S., Kabani, N., & Henkelman, R. M. (2008). High resolution three-dimensional brain atlas using an average magnetic resonance image of 40 adult C57Bl/6J mice. NeuroImage, 42, 60–69.

van Eede, M. C., Scholz, J., Chakravarty, M. M., Henkelman, R. M., & Lerch, J. P. (2013). Mapping registration sensitivity in MR mouse brain images. NeuroImage, 82, 226–236.

Ellegood, J., Anagnostou, E., Babineau, B., Crawley, J., Lin, L., Genestine, M., DiCicco-Bloom, E., Lai, J., Foster, J., Penagarikano, O. et al. (2015). Clustering autism: using neuroanatomical differences in 26 mouse models to gain insight into the heterogeneity. Molecular Psychiatry, 20, 118–125.

Evans A. C. (2013). Networks of anatomical covariance. NeuroImage, 80, 489–504.

French, L., & Pavlidis, P. (2011). Relationships between gene expression and brain wiring in the adult rodent brain. PLOS Computational Biology, 7, e1001049.

Friedel, M., van Eede, M. C., Pipitone, J., Chakravarty, M. M., & Lerch, J. P. (2014). Pydpiper: A flexible toolkit for constructing novel registration pipelines. Frontiers in Neuroinformatics, 8, 67.

Gong, G., He, Y., Chen, Z. J., & Evans, A. C. (2012). Convergence and divergence of thickness correlations with diffusion connections across the human cerebral cortex. NeuroImage, 59, 1239–1248.

Grandjean, J., Zerbi, V., Balsters, J., Wenderoth, N., & Rudin, M. (2017). The structural basis of large-scale functional connectivity in the mouse. The Journal of Neuroscience, (pp. 0438–17).

de Guzman, A. E., Wong, M. D., Gleave, J. A., & Nieman, B. J. (2016). Variations in post-perfusion immersion fixation and storage alter MRI measurements of mouse brain morphometry. NeuroImage, 142, 687–695.

Hänggi, J., Wotruba, D., & Jäncke, L. (2011). Globally altered structural brain network topology in grapheme-color synesthesia. The Journal of Neuroscience, 31, 5816–5828.

He, Y., Chen, Z. J., & Evans, A. C. (2007). Small-world anatomical networks in the human brain revealed by cortical thickness from MRI. Cerebral Cortex, 17, 2407–2419.

Hintiryan, H., Foster, N. N., Bowman, I., Bay, M., Song, M. Y., Gou, L., Yamashita, S., Bienkowski, M. S., Zingg, B., Zhu, M. et al. (2016). The mouse cortico-striatal projectome. Nature Neuroscience, 19, 1100–1114.

Honey, C., Sporns, O., Cammoun, L., Gigandet, X., Thiran, J.-P., Meuli, R., & Hagmann, P. (2009). Predicting human resting-state functional connectivity from structural connectivity. Proceedings of the National Academy of Sciences, 106, 2035–2040.

Jeurissen, B., Leemans, A., Tournier, J.-D., Jones, D. K., & Sijbers, J. (2013). Investigating the prevalence of complex fiber configurations in white matter tissue with diffusion magnetic resonance imaging. Human Brain Mapping, 34, 2747–2766.

Koenigs, M., & Grafman, J. (2009). The functional neuroanatomy of depression: distinct roles for ventromedial and dorsolateral prefrontal cortex. Behavioural Brain Research, 201, 239–243.

Krienen, F. M., Yeo, B. T., Ge, T., Buckner, R. L., & Sherwood, C. C. (2016). Transcriptional profiles of supragranular-enriched genes associate with corticocortical network architecture in the human brain. Proceedings of the National Academy of Sciences, 113, E469–E478.

Lein, E. S., Hawrylycz, M. J., Ao, N., Ayres, M., Bensinger, A., Bernard, A., Boe, A. F., Boguski, M. S., Brockway, K. S., Byrnes, E. J. et al. (2007). Genome-wide atlas of gene expression in the adult mouse brain. Nature, 445, 168–176.

Leow, A. D., Yanovsky, I., Chiang, M.-C., Lee, A. D., Klunder, A. D., Lu, A., Becker, J. T., Davis, S. W., Toga, A. W., & Thompson, P. M. (2007). Statistical properties of Jacobian maps and the realization of unbiased large-deformation nonlinear image registration. IEEE Transactions on Medical Imaging, 26, 822–832.

Lerch, J. P., Sled, J. G., & Henkelman, R. M. (2011a). MRI phenotyping of genetically altered mice. Methods in Molecular Biology, 711, 349–361.

Lerch, J. P., Worsley, K., Shaw, W. P., Greenstein, D. K., Lenroot, R. K., Giedd, J., & Evans, A. C. (2006). Mapping anatomical correlations across cerebral cortex (MACACC) using cortical thickness from MRI. NeuroImage, 31, 993–1003.

Lerch, J. P., Yiu, A. P., Martinez-Canabal, A., Pekar, T., Bohbot, V. D., Frankland, P. W., Henkelman, R. M., Josselyn, S. A., & Sled, J. G. (2011b). Maze training in mice induces MRI-detectable brain shape changes specific to the type of learning. NeuroImage, 54, 2086–2095.

Loken, C., Gruner, D., Groer, L., Peltier, R., Bunn, N., Craig, M., Henriques, T., Dempsey, J., Yu, C.-H., Chen, J. et al. (2010). SciNet: lessons learned from building a power-efficient top-20 system and data centre. Journal of Physics: Conference Series, 256, 012026.

Maguire, E. A., Gadian, D. G., Johnsrude, I. S., Good, C. D., Ashburner, J., Frackowiak, R. S., & Frith, C. D. (2000). Navigation-related structural change in the hippocampi of taxi drivers. Proceedings of the National Academy of Sciences, 97, 4398–4403.

McFadden D. (1974). Conditional logit analysis of qualitative choice behavior. In P. Zarembka (Ed.), Frontiers in Econometrics chapter 4. (pp. 105–142). New York: Academic Press.

Mills, B. D., Grayson, D., Shunmugavel, A., Miranda-Dominguez, O., Feczko, E., Earl, E., Neve, K., & Fair, D. (2017). Correlated gene expression and anatomical communication support synchronized brain activity in the mouse functional connectome. bioRxiv, eprint. doi: 10.1101/167304.

Murphy, K., Birn, R. M., Handwerker, D. A., Jones, T. B., & Bandettini, P. A. (2009). The impact of global signal regression on resting state correlations: are anti-correlated networks introduced? NeuroImage, 44, 893–905.

Oh, S. W., Harris, J. A., Ng, L., Winslow, B., Cain, N., Mihalas, S., Wang, Q., Lau, C., Kuan, L., Henry, A. M. et al. (2014). A mesoscale connectome of the mouse brain. Nature, 508, 207–214.

Pagani, M., Bifone, A., & Gozzi, A. (2016). Structural covariance networks in the mouse brain. NeuroImage, 129, 55–63.

Pezawas, L., Meyer-Lindenberg, A., Goldman, A., Verchinski, B., Chen, G., Kolachana, B., Egan, M., Mattay, V., Hariri, A., & Weinberger, D. (2008). Evidence of biologic epistasis between BDNF and SLC6A4 and implications for depression. Molecular Psychiatry, 13, 709–716.

Plachez, C., & Richards, L. J. (2005). Mechanisms of axon guidance in the developing nervous system. Current Topics in Developmental Biology, 69, 267–346.

Raznahan, A., Lerch, J. P., Lee, N., Greenstein, D., Wallace, G. L., Stockman, M., Clasen, L., Shaw, P. W., & Giedd, J. N. (2011). Patterns of coordinated anatomical change inhuman cortical development: a longitudinal neuroimaging study of maturational coupling. Neuron, 72, 873–884.

Reid, A. T., Lewis, J., Bezgin, G., Khundrakpam, B., Eickhoff, S. B., McIntosh, A. R., Bellec, P., & Evans, A. C. (2016). A cross-modal, cross-species comparison of connectivity measures in the primate brain. NeuroImage, 125, 311–331.

Richiardi, J., Altmann, A., Milazzo, A.-C., Chang, C., Chakravarty, M. M., Banaschewski, T., Barker, G. J., Bokde, A. L., Bromberg, U., Büchel, C. et al. (2015). Correlated gene expression supports synchronous activity in brain networks. Science, 348, 1241–1244.

Rimol, L. M., Panizzon, M. S., Fennema-Notestine, C., Eyler, L. T., Fischl, B., Franz, C. E., Hagler, D. J., Lyons, M. J., Neale, M. C., Pacheco, J. et al. (2010). Cortical thickness is influenced by regionally specific genetic factors. Biological Psychiatry, 67, 493–499.

Romero Garcia, R., Whitaker, K., Vasa, F., Seidlitz, J., Shinn, M., Fonagy, P., Dolan, R., Jones, P., Goodyer, I., Bullmore, E., & Vertes, P. (2017). Structural covariance networks are coupled to expression of genes enriched in supragranular layers of the human cortex. bioRxiv, eprint. doi: 10.1101/163758.

Scannell, J. W., Blakemore, C., & Young, M. P. (1995). Analysis of connectivity in the cat cerebral cortex. The Journal of Neuroscience, 15, 1463–1483.

Schmitt, J., Lenroot, R., Wallace, G., Ordaz, S., Taylor, K., Kabani, N., Greenstein, D., Lerch, J., Kendler, K., Neale, M. et al. (2008). Identification of genetically mediated cortical networks: a multivariate study of pediatric twins and siblings. Cerebral Cortex, 18, 1737–1747.

Schmitt, J. E., Yi, J., Calkins, M. E., Ruparel, K., Roalf, D. R., Cassidy, A., Souders, M. C., Satterthwaite, T. D., McDonald-McGinn, D. M., Zackai, E. H. et al. (2016). Disrupted anatomic networks in the 22q11.2 deletion syndrome. NeuroImage: Clinical, 12, 420428.

Segall, J. M., Allen, E. A., Jung, R. E., Erhardt, E. B., Arja, S. K., Kiehl, K. A., & Calhoun, V. D. (2012). Correspondence between structure and function in the human brain at rest. Frontiers in Neuroinformatics, 6, 10.

Sethian J. A. (1996). A fast marching level set method for monotonically advancing fronts. Proceedings of the National Academy of Sciences, 93, 1591–1595.

Shi, F., Yap, P.-T., Gao, W., Lin, W., Gilmore, J. H., & Shen, D. (2012). Altered structural connectivity in neonates at genetic risk for schizophrenia: a combined study using morphological and white matter networks. NeuroImage, 62, 1622–1633.

Steadman, P. E., Ellegood, J., Szulc, K. U., Turnbull, D. H., Joyner, A. L., Henkelman, R. M., & Lerch, J. P. (2014). Genetic effects on cerebellar structure across mouse models of autism using a magnetic resonance imaging atlas. Autism Research, 7, 124–137.

Ullmann, J. F., Watson, C., Janke, A. L., Kurniawan, N. D., & Reutens, D. C. (2013). A segmentation protocol and MRI atlas of the C57BL/6J mouse neocortex. NeuroImage, 78, 196–203.

Valk, S. L., Di Martino, A., Milham, M. P., & Bernhardt, B. C. (2015). Multicenter mapping of structural network alterations in autism. Human Brain Mapping, 36, 2364–2373.

Vasa, F., Seidlitz, J., Romero-Garcia, R., Whitaker, K. J., Rosenthal, G., Vertes, P. E., Shinn, M., Alexander-Bloch, A., Fonagy, P., Dolan, R., Jones, P., Goodyer, I., The NSPN Consortium, Sporns, O., & Bullmore, E. T. (2017). Adolescent tuning of association cortex in human structural brain networks. bioRxiv, eprint. doi:10.1101/126920.

Vértes, P. E., Rittman, T., Whitaker, K. J., Romero-Garcia, R., Váša, F., Kitzbichler, M. G., Wagstyl, K., Fonagy, P., Dolan, R. J., Jones, P. B. et al. (2016). Gene transcription profiles associated with inter-modular hubs and connection distance in human functional magnetic resonance imaging networks. Philosophical Transactions of the Royal Society B, 371, 20150362.

Voss, P., & Zatorre, R. J. (2015). Early visual deprivation changes cortical anatomical covariance in dorsal-stream structures. NeuroImage, 108, 194–202.

Wheeler, A. L., Chakravarty, M. M., Lerch, J. P., Pipitone, J., Daskalakis, Z. J., Rajji, T. K., Mulsant, B. H., & Voineskos, A. N. (2014). Disrupted prefrontal interhemispheric structural coupling in schizophrenia related to working memory performance. Schizophrenia Bulletin, 40, 914–924.

Yasuda, C. L., Chen, Z., Beltramini, G. C., Coan, A. C., Morita, M. E., Kubota, B., Bergo, F., Beaulieu, C., Cendes, F., & Gross, D. W. (2015). Aberrant topological patterns of brain structural network in temporal lobe epilepsy. Epilepsia, 56, 1992–2002.

Zielinski, B. A., Anderson, J. S., Froehlich, A. L., Prigge, M. B., Nielsen, J. A., Cooperrider, J. R., Cariello, A. N., Fletcher, P. T., Alexander, A. L., Lange, N. et al. (2012). scMRI reveals large-scale brain network abnormalities in autism. PLOS ONE, 7, e49172.

